# Phylogenetic Diversity Across the Complete Tree of Life

**DOI:** 10.1101/2025.08.12.669825

**Authors:** Jialiang Guo, Emily Jane McTavish, James Rosindell

**Affiliations:** Peking University; University of California, Merced; Imperial College London

**Keywords:** Conservation, Evolutionary Distinctiveness, Open Tree of Life, Phylogenetic Diversity

## Abstract

In the face of rapid biodiversity loss, many approaches have been developed to measure biodiversity in ways that go beyond species richness. One prominent example is Phylogenetic Diversity (PD), which, on a dated phylogenetic tree, measures evolutionary history by summing the branch lengths measured in time required to connect a set of species. PD may also capture other biodiversity measures by proxy such as the richness of biological features and their potential benefits for humanity, known as ‘future options’. The total global PD is known for some well-studied groups, such as most vertebrates, but PD estimates are lacking for the majority of the tree of life. Here, we characterize the distribution of PD across the complete tree of life with over 2.2 million species. To do this we use data from the Open Tree of Life and a smoothing method to interpolate between nodes without date information. We estimate that the PD represented by all described species together is between 30 and 33 trillion years (95% confidence interval). We characterize the distribution of evolutionary distinctiveness, a measure of the fair share of PD captured by individual species, across all life and within selected clades. This Evolutionary Distinctiveness metric has previously been used as the basis for conservation prioritization schemes such as EDGE (Evolutionary Distinct and Globally Endangered) which synthesizes phylogenetic tree data with extinction risk data from the IUCN Red List of threatened species. We use our results to estimate EDGE scores for over 130,000 species. We also estimate the amount of threatened phylogenetic diversity across the complete tree of life. We hope this work will pave the way for more complete and automated analyses of PD, Evolutionary Distinctiveness, threatened evolutionary history, and EDGE scores across the complete tree of life.

## 1. Introduction

We are currently experiencing a massive decline in global biodiversity, the variety of life on earth. Anthropogenic activities such as livestock grazing, urbanization, and emission of greenhouse gasses are reshaping the land, driving what is being described as the sixth mass extinction on our planet^1–3^. This decline threatens the stability of many ecosystem services which are crucial for the quality of human life now and in the future^4^. There are over 2.2 million described species of life on earth, but species richness is not the only dimension of biodiversity and its decline. Other measures of biodiversity such as trait and functional diversity use details about individual species that are currently unknown for the vast majority of described species. A complete tree of life (phylogeny) describing the evolutionary connections between species is available however, and could be used to measure deeper aspects of biodiversity at a global scale that are not captured by species richness.

The most commonly used measure of biodiversity based on data from the tree of life is Phylogenetic Diversity (PD), which measures the evolutionary history of a group of species. PD is calculated by summing the phylogenetic branch lengths required to span a given group of species^5^. This biodiversity measure can be regarded as a proxy for other biodiversity measures, especially feature diversity (the diversity of biological features or traits in the most general sense that can be detected in their structure or biochemistry). PD may potentially also capture ‘future options’ for humanity: features of species with potential utility for humans, which are currently unknown, but which may manifest in the future^6–9^.

The importance of preserving PD has been increasingly recognized, the Intergovernmental Science-policy Platform for Biodiversity and Ecosystem Services (IPBES) has adopted PD as an indicator of the overall capacity of biodiversity to support a good quality of life into the future^6^. This indicator, along with the related ‘EDGE’ (Evolutionary Distinct and Globally Endangered) Index^10^, have been included as indicators for the Convention on Biological Diversity’s draft post-2020 Global Biodiversity Framework^11^. The IUCN has also established a Phylogenetic Diversity Task Force to provide expertise on the inclusion of PD in conservation strategies for practitioners, decision-makers, and the public^12^. Another important measure related to PD is the Evolutionary Distinctiveness (ED) of a species, also known as ‘fair proportion’^10,13^. This measure represents a fair share of the total PD of a given clade that is attributed to each individual one of its species. ED can therefore be used to measure the contribution made by a specific individual species to phylogenetic diversity of a larger clade^13^. In the calculation of ED for a given species on a phylogeny, we consider separately all its ancestral branches back to the root. For each branch, the branch length, which comprises a contribution to PD, is divided equally (fairly) between all its descendent extant species (tips). The total PD across the complete tree of life is equal to the sum of ED across all species. We can, therefore, use ED calculation or approximation as a way to find total PD through summation. However, most often, PD and ED are calculated together using a phylogenetic tree with branch lengths.

One application of PD and ED is in conservation prioritization. Many studies have used the idea of preserving PD in preference to metrics based only on species richness and to augment endangerment data as a way to guide species-based conservation priorities. For example, the EDGE score of a species is calculated by combining its ED score and the risk of extinction provided by the IUCN Red List. The EDGE metric enables researchers to identify the most evolutionarily unique and threatened species as potential priorities^10,14,15^ in the context of limited resources for practical conservation^16^. This ranking has been done for at least 7 clades including gymnosperms^14^, corals^17^, amphibians^18^, chondrichthyes^19^, reptiles^20^, mammals and birds^10,21–26^. However, most of the tree of life has not been explored in this way.

Here we make use of data from the Open Tree of Life (OpenTree)^27^ and OneZoom tree of life explorer^28,29^ to study PD and EDGE scores across all described species of life. The OpenTree project builds and maintains a comprehensive tree of life by integrating published phylogenies with taxonomic information^30^. This synthetic tree is updated using automated pipelines^31^ and contains more than 2.3 million species with each species represented by a unique identifier in the ‘Open Tree Taxonomy’ (OTT). We use date estimates from published phylogenetic studies in the Open Tree data store mapped to internal nodes on the OpenTree synthetic phylogeny^30^. OneZoom synthesizes the OpenTree phylogeny data with a curated backbone and other published phylogenies to provide a way to visualize the tree of life for broader audiences, and combines the phylogenetic data with the risk of extinction provided by the IUCN Red List. Together, these tools provide an opportunity to study phylogenetic diversity and evolutionary distinctiveness at the scale of all described life on earth.

We explore how the ED scores are distributed between species across the complete tree of life, focussing on selected major taxonomic groups. We estimate the total PD, and proportion of threatened PD for different clades of species, and contrast this to species richness. Finally, we estimate EDGE scores for all species that are evaluated by the IUCN red list of threatened species^32^. Whilst EDGE-based studies of increasingly large clades have been done before^10,14,15^, to the best of our knowledge, this is the first time that all described species with IUCN Red List data have been given EDGE scores.

We chose to include well known groups to aid comparison with other studies, as well as some more obscure ones based on them being monophyletic groups with distinctive positions in the tree of life. This meant accepting a known bias in existing work with our choice of taxa, whilst also including all life and selected understudied areas. Crucially, the bias in our study only manifests in how finely we subdivide taxa for comparison to previous work and inevitably in the availability of input data. We do not add biases of our own onto this backdrop and our scope of work includes all described species (see supplementary figure S1).

## 2. Results

The median ED score across all described species of life is 6.36 Myr (2.5^th^ and 97.5^th^ centiles are 0.23 - 65.74 Myr). We analyzed all described Eukaryotic life across 23 selected monophyletic clades (see supplementary figure S1). We also produced a table of ED scores across the rest of the tree of life (see supplementary table S1) Species in the TSAR (Telonemia, Stramenopile, Alveolates and Rhizaria) clade have a median ED score of 33.97Myr in the context of the complete tree of life. For Mollusca species median ED is 24.45 Myr and for the Cyclostomata clade median ED is 20.18 Myr. Among the 9 selected Vertebrata clades, there is relative consistency between the median ED scores of Actinopterygii (7.84 Myr), Amphibia (8.33 Myr) and Mammalia (8.21 Myr) (see figure 1).

**Figure 1.**
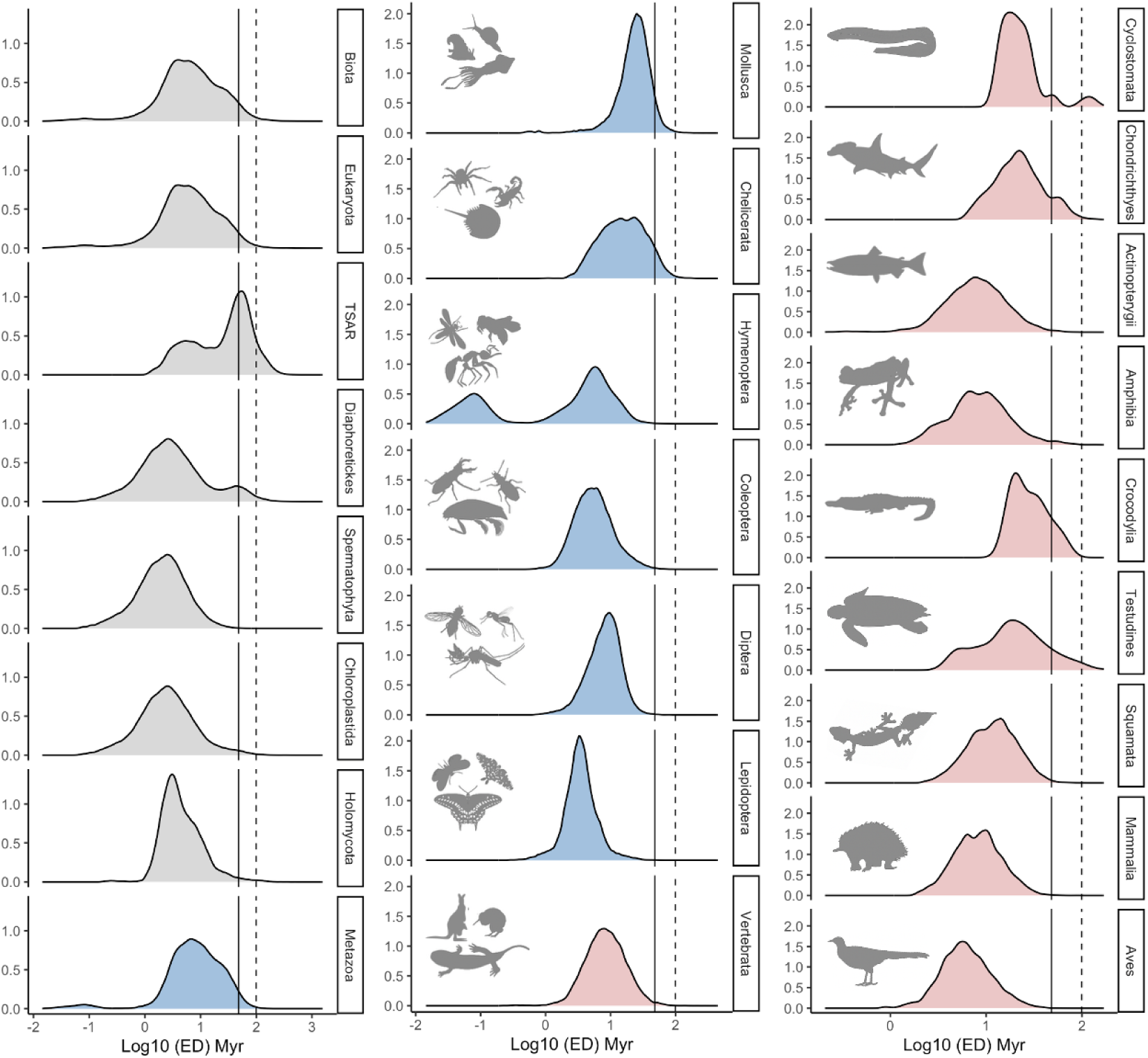
The distribution of ED scores across all species on log-scale, shown separately for selected monophyletic clades of species. Each species’ ED score incorporated into these distributions represents the median of 1000 replicates in a jackknife analysis - see methods. The 10 selected vertebrata clades are shown in red, the 7 selected metazoan clades (other than vertebrata) are shown in blue, and the 6 selected clades from Eukaryota (other than metazoa) are shown in grey. All clades are labelled with text and the clade corresponding to all described life (Biota) is shown in the first panel at the top left. The solid vertical line in each panel represents the 95% quantile of log ED score across all described life, which is 1.681, and the dashed line represents the 99% quantile, which is 1.997. Silhouette image credits are given in the paper acknowledgements.

We observe a bimodal distribution of ED scores in the TSAR, Diaphoretickes and Hymenoptera clades (figure. 1), however, large subclades of these groups usually have a single mode. For example, in the TSAR clade, the Rhizaria and Alveolata subclades, account for 61% of the species richness and have a single mode in their ED distribution. (Supplementary figure S3).

The ED scores across all life appear log-normal but only superficially as they fail the test of log-normality (Biota, Eukaryota, TSAR, Diaphoretickes, Spermatophyta, Chloroplastida, Holomycota, Metazoa, Mollusca, Chelicerata, Hymenoptera, Coleoptera, Diptera, Lepidoptera, Vertebrata, Actinopterygii, Aves: p-value < 2.2e-16, Cyclostomata: p = 6.853e-13, Chondrichthyes: p = 3.477e-05,: p < 2.2e-16, Amphibia: p = 2.837e-13, Testudines: p = 0.03005, Squamata: p = 8.316e-14, Mammalia: p = 0.002891, Anderson-Darling normality test, Crocodilia: p = 0.04708, Shapiro-Wilk normality test). The departure from log-normality is primarily in the tails of the distribution as can be seen from a Quantile-Quantile (Q-Q) plot (Supplementary figure S2).

We identified the number and proportion of species within each selected monophyletic clade that fall into the top 1% (99.3 Myr), top 5% (48.01 Myr) and top 50% (6.36 Myr) of Evolutionary Distinctiveness (ED) across all described life. According to our randomisation test, among all selected monophyletic clades, TSAR, Diaphoretickes, Cyclostomata and Testudines have significantly more of life’s most evolutionarily distinct species than would be expected by chance if evolutionary distinctiveness scores were all randomly shuffled across all species. The other selected clades contain fewer of life’s most evolutionary distinct species than would be expected by chance (figure. 2)

**Figure 2.**
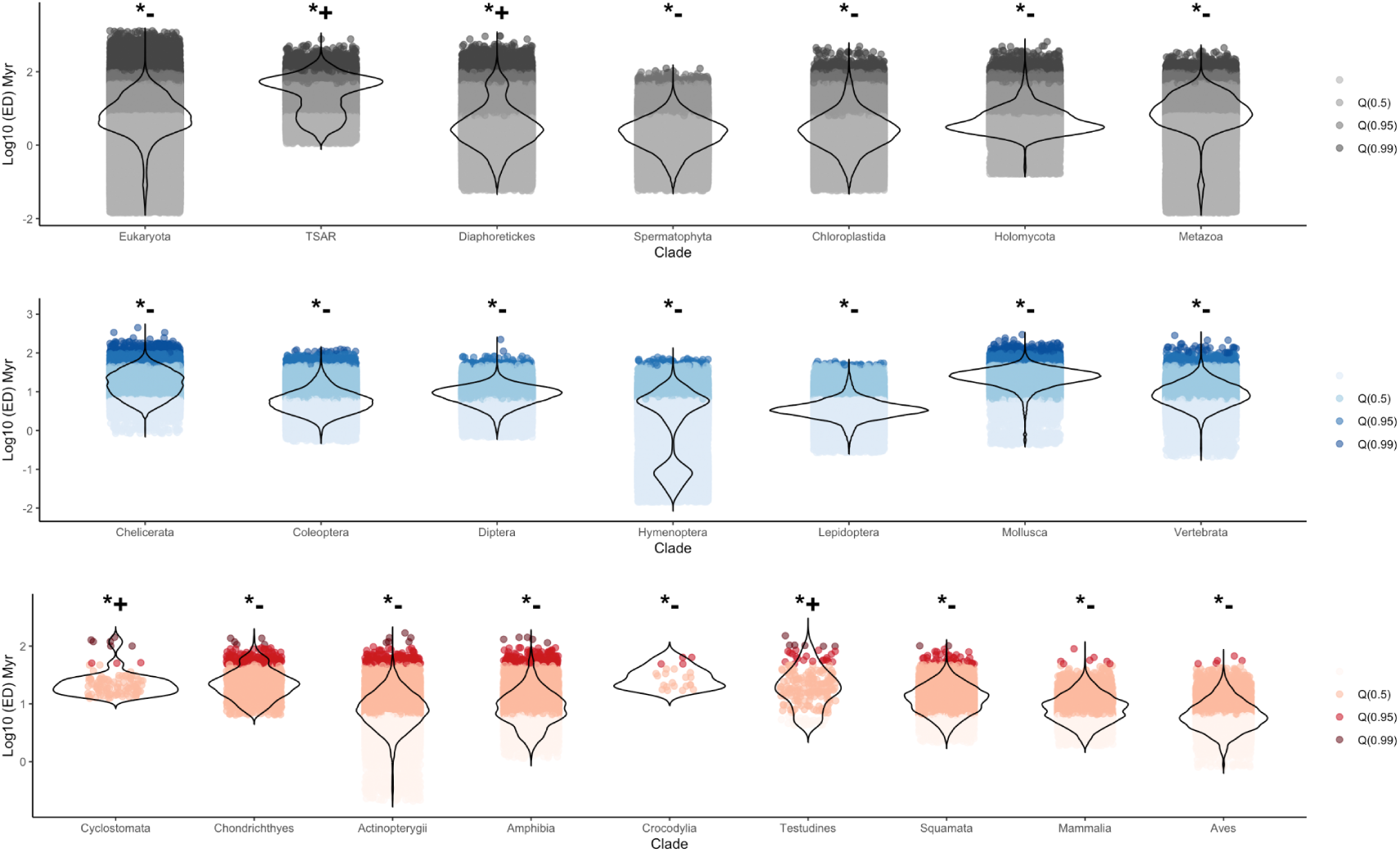
The proportion of high ED species among selected clades. The violin plot shows the distribution of ED scores in different clades and coloured points show species whose ED score is above the 99% quantile, 95% quantile and 50% quantile (median) of ED score of all described life on earth. The \**+* indicates this clade has significantly larger proportion of evolutionary distinctive (95% quantile in the context of all life) species than expected by chance from a random shuffling of ED among species, while the *- indicates this clade has a significantly smaller proportion of evolutionary distinctive species than expected by chance.

The median total unique PD (across 1000 replicates of a jackknife analysis) for all life calculated in our study was 31.36 trillion years with values ranging from 30 to 33 trillion years across all replicates. Our results show that 14 out of the 23 selected clades have robust estimates of total unique PD with a jackknife analysis relative 95% confidence interval width of no more than 84% of the median PD. In contrast, the Cyclostomata, Chondrichthyes, Crocodylia, Lepidoptera, Diptera, Hymenoptera, Mollusca, Holomycota and TSAR clades showed a higher degree of uncertainty with relative 95% confidence interval widths of 485%, 127%, 216%, 134%, 169%, 190%, 139%, 130% and 143% respectively (shown as percentages of the median estimated PD) (see figure 3). PD scores across the complete set of monophyletic higher taxa are available as supplementary information (see supplementary table S2).

**Figure 3.**
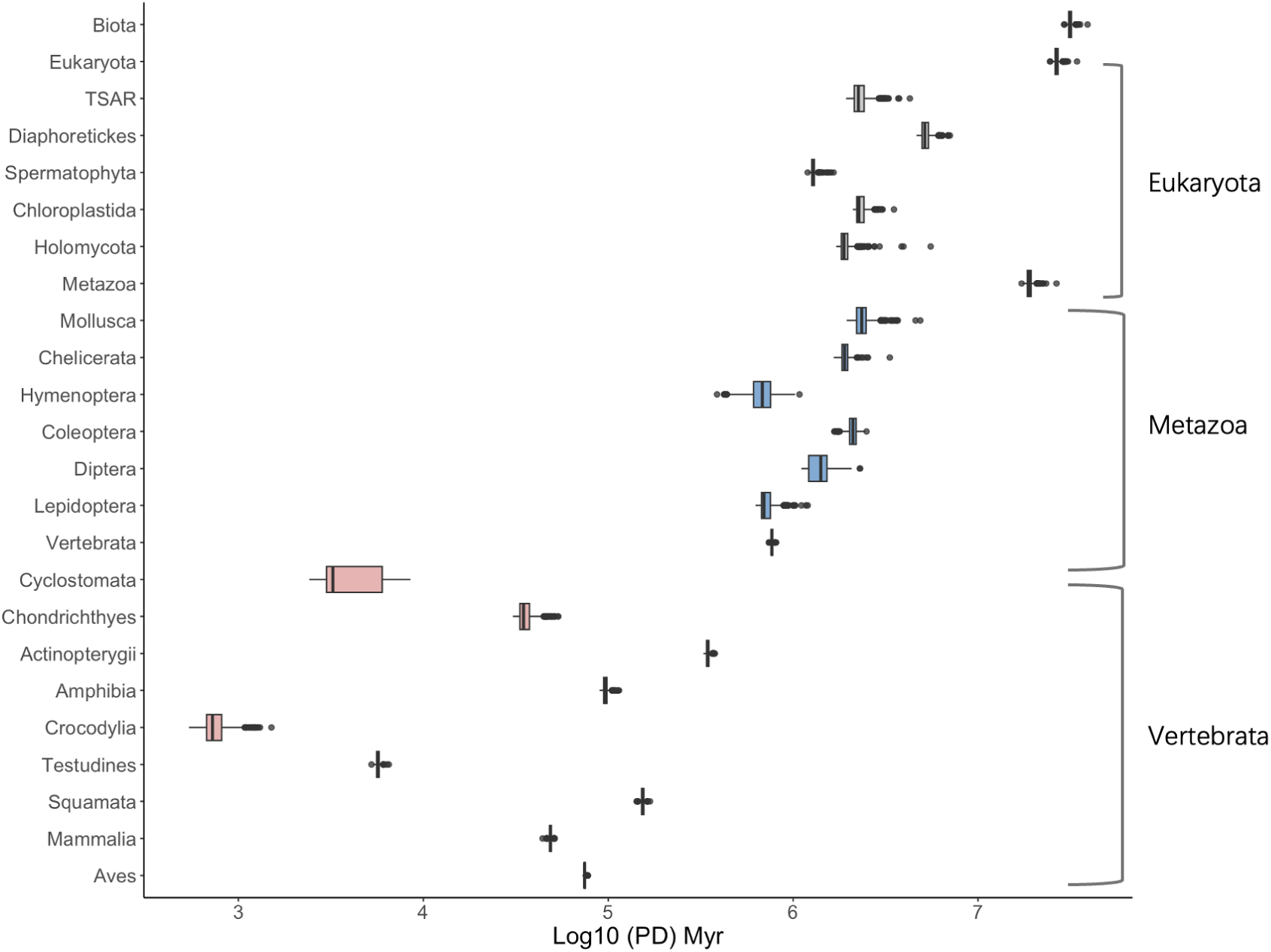
The distribution of total unique PD scores across selected monophyletic clades. Boxplots with median (solid line), 25th and 75th percentiles (box edges), and 5th and 95th centiles (whiskers). Colors follow the same scheme as figures 1 and 2. The total unique PD values shown do not include a stem back to the origin of life to enable comparability and thus are slightly smaller than the sum of the ED scores of all species in the clade because the sum of ED scores would only include a fair share of the stem.

The ratio of PD to species richness varied between 3.47 and 45.99 Myr per species among our selected taxa. Seven taxa contribute disproportionately much to the total unique PD given their total number of species: TSAR, Mollusca, Chelicerata, Cyclostomata, Chondrichthyes, Crocodylia and Testudines all of which had a ratio greater than 20 Myr per species (see Table 1 for all numbers). Because our groups are considered in the context of all life we could include all the branches back to the origin of life as part of the PD for each group. We have chosen not to do this and so we are technically reporting ‘unique’ PD, that is the PD that is not covered by any species outside the focal group. In contrast, The Spermatophyta, Lepidoptera, Hymenoptera and Aves clades represent low unique PD for their species richness, all below 7.5 Myr per species (see Table 1). Among all the selected Eukaryota clades, Metazoa has the highest median total unique PD (18.9 Tyr - trillion years), which can be attributed to the significantly higher species richness compared with other Eukaryota clades. Actinopterygii, is the Vertebrata clade that has both the highest species richness and the highest median total unique PD (345,221 Myr).

**Table 1.**
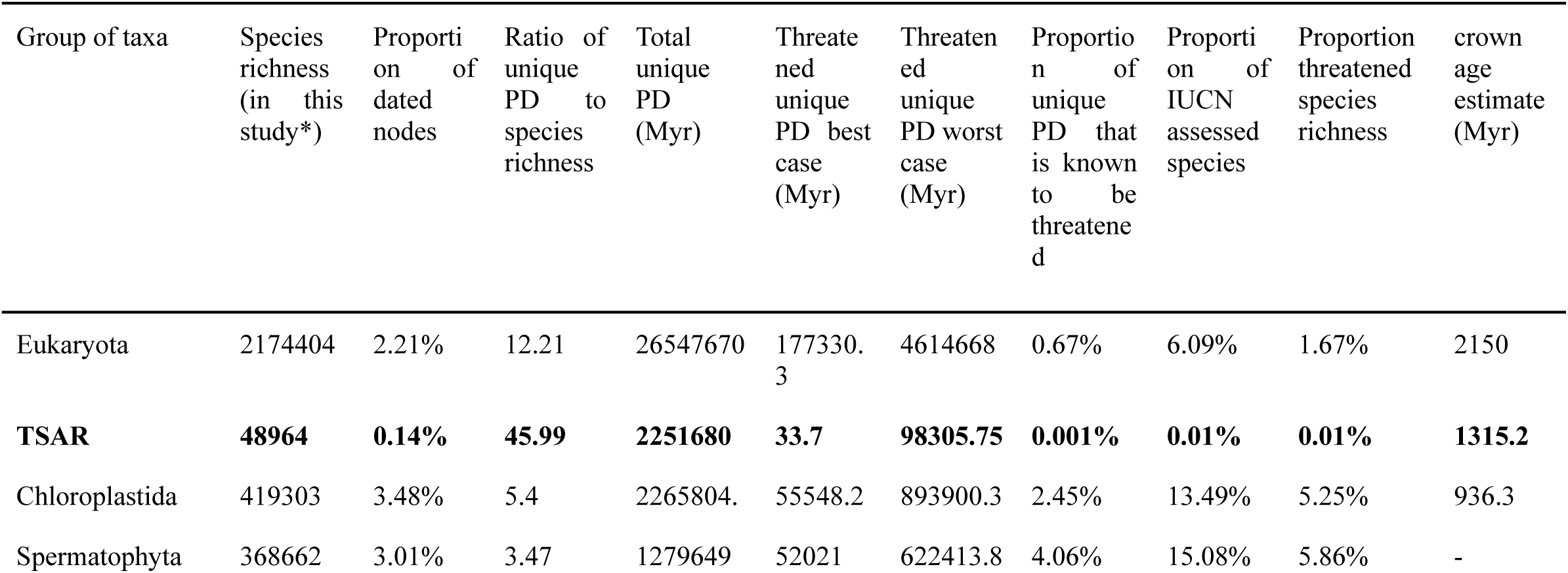
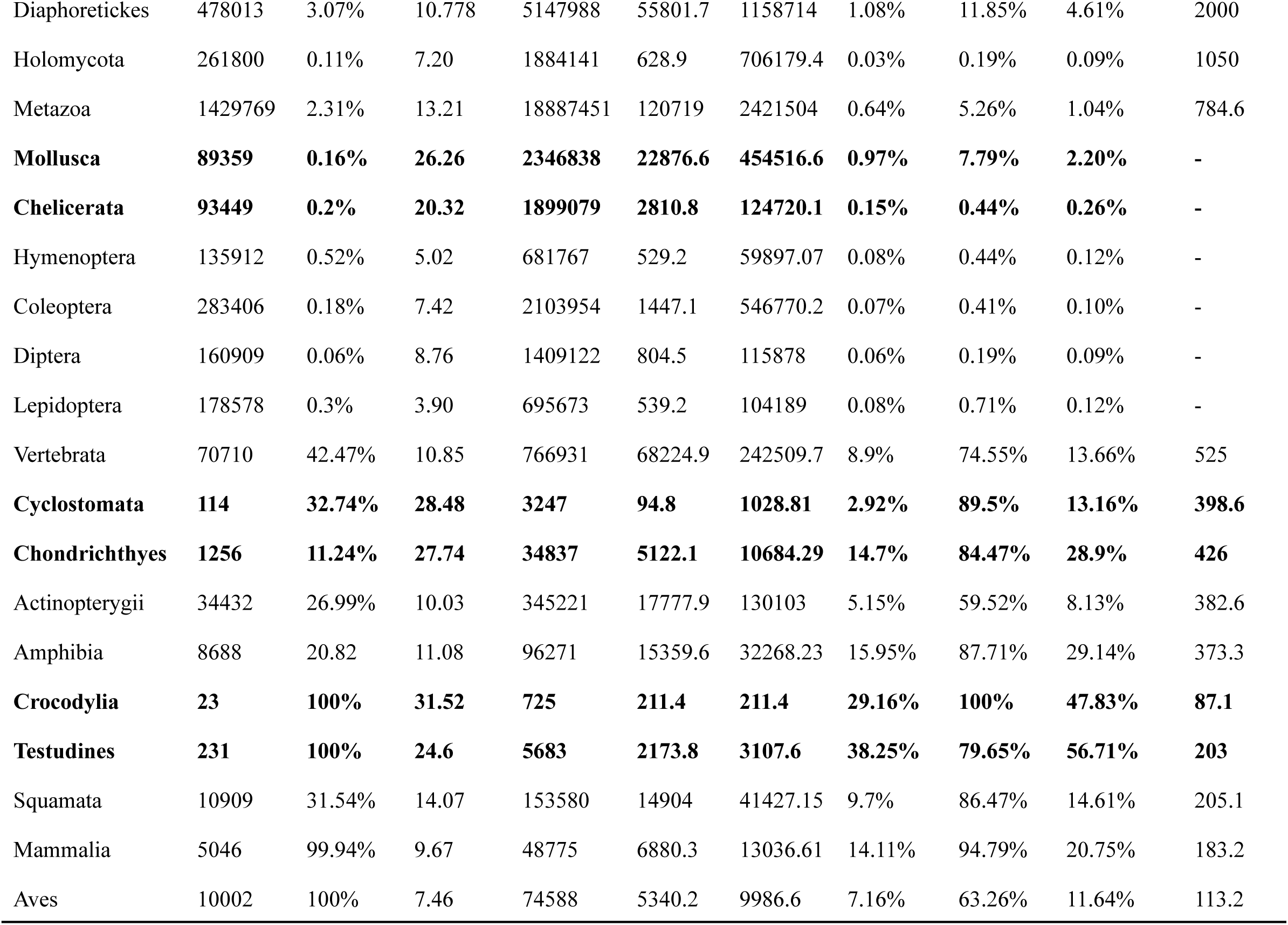
The relationship between species richness and total unique PD. The columns of the table have the following meanings: i) Group of taxa - this is a monophyletic group; ii) Species richness - a count of the total number of species in the group - this is a total of the species in our study for comparison and context, it should not be taken as a definitive richness value for the listed groups; iii) Proportion of dated nodes - based on available data from OpenTree combined with OneZoom which incorporates some fully dated sub trees iv) Ratio of PD to species richness in Myr per species; v) total unique PD - this excludes the stem back to the origin of life but includes the shorter stem which connects to the group to its closest related outgroup vi) Threatened unique PD best case - a sum of branches within the clade where all descendants are threatened (CR, EN or VU in the red list, DD and NE species were not considered threatened here); vii) Threatened unique PD worst case as for vi) except now DD and NE species are considered threatened; viii) proportion of unique PD that is known to be threatened - based on column vii) as a proportion of column v) and expressed as a percentage; ix) proportion of IUCN assessed species - the proportion of species richness that is covered by a valid IUCN assessment i.e. excluding those which are not evaluated. x) Proportion of species richness known to be threatened (CR, EN or VU) xi) crown age in Myr - crown age estimates are only included where we had dates in source data, they were not interpolated. The 7 clades highlighted in boldface have a ratio of total unique PD / species richness of at least 20 Myr per species. See methods section for further details.

| Group of taxa | Species richness (in this study*) | Proportion of dated nodes | Ratio of unique PD to species richness | Total unique PD (Myr) | Threatened unique PD best case (Myr) | Threatened unique PD worst case (Myr) | Proportion of unique PD that is known to be threatened | Proportion of IUCN assessed species | Proportion of threatened species richness | crown age estimate (Myr) |
| --- | --- | --- | --- | --- | --- | --- | --- | --- | --- | --- |
| Eukaryota | 2174404 | 2.21% | 12.21 | 26547670 | 177330.3 | 4614668 | 0.67% | 6.09% | 1.67% | 2150 |
| **TSAR** | **48964** | **0.14%** | **45.99** | **2251680** | **33.7** | **98305.75** | **0.001%** | **0.01%** | **0.01%** | **1315.2** |
| Chloroplastida | 419303 | 3.48% | 5.4 | 2265804. | 55548.2 | 893900.3 | 2.45% | 13.49% | 5.25% | 936.3 |
| Spermatophyta | 368662 | 3.01% | 3.47 | 1279649 | 52021 | 622413.8 | 4.06% | 15.08% | 5.86% | - |
| Diaphoretickes | 478013 | 3.07% | 10.778 | 5147988 | 55801.7 | 1158714 | 1.08% | 11.85% | 4.61% | 2000 |
| Holomycota | 261800 | 0.11% | 7.20 | 1884141 | 628.9 | 706179.4 | 0.03% | 0.19% | 0.09% | 1050 |
| Metazoa | 1429769 | 2.31% | 13.21 | 18887451 | 120719 | 2421504 | 0.64% | 5.26% | 1.04% | 784.6 |
| **Mollusca** | **89359** | **0.16%** | **26.26** | **2346838** | **22876.6** | **454516.6** | **0.97%** | **7.79%** | **2.20%** | - |
| **Chelicerata** | **93449** | **0.2%** | **20.32** | **1899079** | **2810.8** | **124720.1** | **0.15%** | **0.44%** | **0.26%** | - |
| Hymenoptera | 135912 | 0.52% | 5.02 | 681767 | 529.2 | 59897.07 | 0.08% | 0.44% | 0.12% | - |
| Coleoptera | 283406 | 0.18% | 7.42 | 2103954 | 1447.1 | 546770.2 | 0.07% | 0.41% | 0.10% | - |
| Diptera | 160909 | 0.06% | 8.76 | 1409122 | 804.5 | 115878 | 0.06% | 0.19% | 0.09% | - |
| Lepidoptera | 178578 | 0.3% | 3.90 | 695673 | 539.2 | 104189 | 0.08% | 0.71% | 0.12% | - |
| Vertebrata | 70710 | 42.47% | 10.85 | 766931 | 68224.9 | 242509.7 | 8.9% | 74.55% | 13.66% | 525 |
| **Cyclostomata** | **114** | **32.74%** | **28.48** | **3247** | **94.8** | **1028.81** | **2.92%** | **89.5%** | **13.16%** | **398.6** |
| **Chondrichthyes** | **1256** | **11.24%** | **27.74** | **34837** | **5122.1** | **10684.29** | **14.7%** | **84.47%** | **28.9%** | **426** |
| Actinopterygii | 34432 | 26.99% | 10.03 | 345221 | 17777.9 | 130103 | 5.15% | 59.52% | 8.13% | 382.6 |
| Amphibia | 8688 | 20.82 | 11.08 | 96271 | 15359.6 | 32268.23 | 15.95% | 87.71% | 29.14% | 373.3 |
| **Crocodylia** | **23** | **100%** | **31.52** | **725** | **211.4** | **211.4** | **29.16%** | **100%** | **47.83%** | **87.1** |
| **Testudines** | **231** | **100%** | **24.6** | **5683** | **2173.8** | **3107.6** | **38.25%** | **79.65%** | **56.71%** | **203** |
| Squamata | 10909 | 31.54% | 14.07 | 153580 | 14904 | 41427.15 | 9.7% | 86.47% | 14.61% | 205.1 |
| Mammalia | 5046 | 99.94% | 9.67 | 48775 | 6880.3 | 13036.61 | 14.11% | 94.79% | 20.75% | 183.2 |
| Aves | 10002 | 100% | 7.46 | 74588 | 5340.2 | 9986.6 | 7.16% | 63.26% | 11.64% | 113.2 |

Out of the clades we considered, Testudines has the highest proportion of known threatened PD (38.7%, 2.2 giga-years, Gy). The least speciose clade, Crocodylia, has the second largest proportion of known threatened PD (29.16%, 0.2 Gy). Amphibians have the third largest proportion of known threatened PD (15.95%, 15.4 Gy) (Table 1). Actinopterygii, the largest vertebrate clade both in terms of species richness and PD, has 17.8 billion years of known threatened PD, which accounts for 5.15% of its total unique PD. The vertebrate clade with the second largest PD was Squamata which has 14.9 Gy of known threatened PD, which accounts for 9.7% of its total unique PD (Table 1).

We calculated EDGE scores for those species that have an IUCN Red List assessment and are aligned to the Open Tree Taxonomy by OneZoom. We ranked the EDGE scores in descending order as a basis for assessing possible conservation priorities. In the top 20 EDGE species there were 3 Reptilia, 3 Copepoda, 3 Amphibia, 1 Coelacanthimorpha, 1 Polypodiopsida, 1 Dipnoi, 1 Florideophyceae, 1 Actinopterygii, 1 Bivalvia, 1 Cycadopsida, 1 Malacostraca, 1 Diplopoda, 1 Marchantiopsida and 1 Gastropoda, no Aves or Mammals were in the top 20 despite a much larger proportion of those groups having Red List assessments (Figure 4). The list of the top 100 EDGE species identified in this study is available in Supplementary Table S1.

**figure 4.**
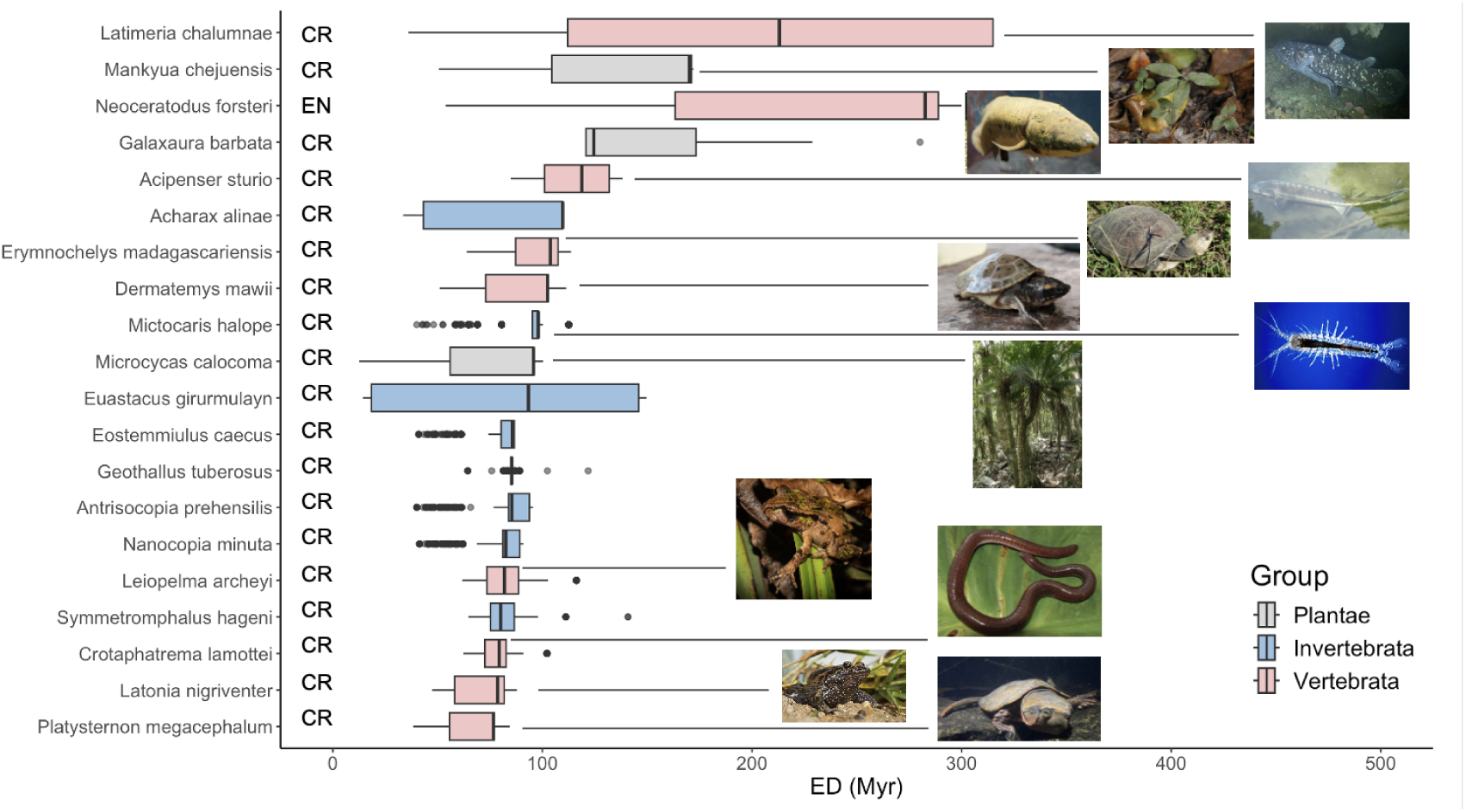
The ED scores of top 20 EDGE species. The images of species which have the highest EDGE score in its class are displayed with their IUCN red list assessment (CR: Critically Endangered; EN: Endangered). The distribution of ED scores for each species here is generated through a jackknife analysis to capture uncertainty in the data. This is done by sampling a subset of the date information and recalculating the result based on extrapolation from only that subset; the resampling was repeated 1000 times (see methods).

## 3. Discussion

The aim of our study was to characterize phylogenetic diversity and evolutionary distinctiveness for all described life on earth. This has only recently been made possible by the arrival of new phylogenetic data adding dates to some nodes on the Open Tree of Life^27,30^. We estimate the total unique PD of all described life to be between 30 and 33 trillion years (95% confidence interval). The numbers are more conservative than previous estimates for large subsets of the complete tree, which suggested 80 trillion years of PD in Metazoa alone. The likely reason is that those estimates assumed a log-linear lineages through time plot and did not incorporate actual tree topology as our study does. Our estimation of PD scores for Actinopterygii, Testudines, Crocodylia, Lepidosaurs and Aves are close to earlier estimations^34,35^ all with a deviation of less than 20% from previously published estimates (Actinopterygii: 4.7%, Testudines: 12.8%, Crocodylia: 18.3%, Lepidosaurs: 14.6% and Aves: 16.9%), only Amphibia had a larger deviation with our study predicting 32.5% less PD compared to previous work (see Supplementary Data S4). This inconsistency could be attributed to the fact that Amphibia had only 20% of nodes having date estimates in our source data. Our estimation of the PD of some clades such as Hymenoptera, Diptera and Cyclostomata (Figure 3) show substantial ranges of uncertainty (Hymenoptera: from 387,449.03 - 1,083,110.3 Myr; Diptera: 1,108,971 - 2,296,654 Myr; Cyclostomata: from 2,417.24 - 8,530.99 Myr). This arises from the combination of a low proportion of dated nodes and resolved nodes (Hymenoptera) and low species richness which accentuates the amount of uncertainty captured by the jackknife analysis process (Cyclostomata). We hope that efforts to generate and collate more dated nodes for large and understudied groups will lead to a natural contraction of these wide uncertainties in future work.

The wide scope of covering all described life, as this study does, inevitably involves caveats. One limitation is the mixed integration of the OneZoom tree of life topology with node dates from the Open Tree of Life. The use of OneZoom meant IUCN Red List data was already aligned with the dataset and the topology benefitted from a degree of further hand curation and direct incorporation of leading studies for certain subclades. OneZoom also resolves polytomies randomly which is necessary to avoid interpreting polytomy uncertainty as increased phylogenetic diversity^36^, however the dataset was built for visualisation and so the polytomy resolution process was only performed once and further uncertainty around it for this analysis is not captured^28^. Another limitation to our dataset is that for well known parts of the tree of life the species richness may not be as accurate as we would like. An example is the testudines which has a low species richness due to having been sourced (though OneZoom) from which has some missing species^37^. Future work should seek to more deeply integrate features of the OneZoom data, Open Tree data and other data sources to produce a distribution of trees capturing more uncertainty. For example, see PhyloNext (https://phylonext.github.io/citations/, DOI: 10.5281/zenodo.7974081) for a promising integration of Open Tree of Life with GBIF. The method for estimating ED and PD itself also has a limitation that a dated tree is never calculated (and is not currently available), estimating ED directly from tree topology and date information means that interior branches without nearby dates would have a different effective length depending on which species we look at for smoothing the dates (see methods). This will have a more substantial effect in parts of the tree with fewer dated nodes and thus more extreme smoothing being required between them. Despite these caveats and future potential for improvement, we expect that at least our first estimate for the PD of all described life will prove useful due to its wide scope and the increasing focus on PD and ED across understudied portions of the tree of life.

Most of our focal clades exhibited unimodal log-normal-like distributions of ED across all their species, the TSAR, Diaphoretickes and Hymenoptera clades instead had bimodal distributions. We hypothesize that the bimodality is caused by heterogeneity among smaller groups within those clades. Many of the clades displaying bimodality can be disaggregated into smaller groups with unimodal distributions of different medians that combine to give an overall bimodal distribution for example a young yet rapidly diversifying clade creates the smaller mode and the remainder of species create the larger mode (Supplementary figure S3). This finding is consistent with observations that patterns of diversification can shift substantively between related monophyletic clades^38,39^. It seems feasible that distributions with more than two modes could occur, but we did not observe this in our groups, possibly because subgroups with extreme values of ED in their species also contain very few species and thus are insignificant in terms of their contribution to the overall ED distribution of the larger clades. A unimodal distribution of ED scores was found in some extremely large clades such as Lepidoptera (178,578 species) and Mollusca (89,359 species). This suggests that even though evolutionary processes are likely heterogeneous at some scale within these clades the heterogeneity is absorbed into a distribution with a single mode, perhaps because the rate shifts affect a large number of small clades each not significant enough to change the qualitative shape of the overall distribution or perhaps because the lack of primary data in those groups restricts our ability to detect rate shifts. Future work may investigate this further by simulating large trees with shifts in speciation and extinction rates and calculating ED distributions on those trees.

Our estimations for the proportion of threatened PD among major tetrapod clades including Amphibian, Mammalia and Aves are largely congruent with an early estimate^34^ (Aves: 7.6 Myr vs 7.46 Myr estimated in our study, Mammalia: 9.6 Myr vs 9.63 Myr estimated in our study, Amphibia: 13.4 Myr vs 11.08 Myr estimated in our study, Chondrichthyes: 30.9 Myr vs 26.18 Myr estimated in our study, Squamata: 12.6 Myr vs 13.75 Myr estimated in our study, all have no more than 20% difference, see Supplementary Data S4), the proportion of threatened PD for Actinopterygii in our study (5.15%) is an underestimate compared to previous work which suggested (12%) of Actinopterygii PD is at risk^35^. The estimation of mean ED scores across different clades in our study are also comparable to previously published estimates (See Supplementary Data S4). However, larger incongruencies can be observed in the comparison of ED scores for certain species, which indicates that lacking phylogenetic information may have a substantial impact on ED score estimation for species embedded in large uncertain regions of the tree (Supplementary figures S7, S8).

Scientists and conservationists alike have long advocated the idea of incorporating phylogenetic diversity into species conservation through use of EDGE-like metrics^40–43^ Here, we combined the ED scores estimated in this study with the extinction risk data provided by the IUCN Red List in order to calculate EDGE scores following (see methods)^10^. Our analysis identified the critically endangered *Latimeria chalumnae* as the top EDGE species. As one of the only two living coelacanth species that are sisters to both lungfish and tetrapods, *L. chalumnae* may help reveal important facts about the adaptation of vertebrate life to land^43^. The coelacanth species also exhibit unique traits such as the intracranial joint and the rostral organ. Given the precarious status of *L. chalumnae* in the wild and its unique features, it is a natural leader on our EDGE list and should be considered a priority for study and for conservation in order to preserve biodiversity in a broad sense^17,45^. The other coelacanth species *Latimeria menadoensis* is vulnerable and thus appears lower down our EDGE ranking, despite its phylogenetic position. Among the top EDGE species in our analysis, the EDGE score estimates for *L. chalumnae*, *N. forsteri*, *M. chejuensis*, *A. sturio*, *E. madagascariensis*, *E. girurmulayn* can be considered reliable because most of their ED score comes from branches of the tree bounded by dated nodes. However, the ED scores of *G. barbata*, *P. chacei*, *M. halope* and *H. huso* are supported by interpolations and random polytomy resolution so whilst they rank highly here they may well rank lower in future analyses where there is less uncertainty (see Supplementary figure S8). The original EDGE metric we use in our study is not the only metric of interest in its class. For example, Heighted EDGE (HEDGE) is a matrix to quantify how much a species is likely to contribute future biodiversity in terms of PD based on potential extinctions^43^. EDGE2 is another more recent advance that incorporates the core principles of HEDGE together with a protocol for its practical use. One advantage of EDGE2 is that the GE component corresponding to endangerment can never be equal to zero, thus it acknowledges that all species have at least some small amount of risk. Future work should attempt to apply EDGE2 to the complete tree of life but that is not possible with our approach because EDGE2 requires calculations on a dated tree with extinction risk data. Here we never dated nodes individually and instead estimated ED directly from the incomplete data. The biggest challenge with expanding to EDGE2 and other measures will come from the dependency on extinction risk for related species because, for most clades in the tree of life, extinction risk information is extremely sparse.

According to our results, the TSAR clade has the highest proportion of species whose ED score is higher than 95% of all described species (35.8%, 17545 species among 48964 TSAR species - see figure 2). Although the TSAR clade accounts for only 2.25% of eukaryotic species richness, this clade contributes 7.2% of the eukaryotic phylogenetic diversity with an average of 45.99 Myr of ED across species (see supplementary figure S5). This is about 5 times higher than the average ED for mammals (9.67 Myr) and about 6 times higher than the average ED for birds (7.47 Myr). Under the assumption that PD does indeed capture important elements of biodiversity such as future options for humanity, this could suggest that species within the TSAR clade should be given some attention for conservation. Indeed, many TSAR species are microorganisms and there are advocates for conserving the biodiversity represented by microorganisms^46,47^. It is possible that much of biodiversity loss, and PD loss is occurring in microorganisms invisible to the human eye. TSAR falls outside the mainstream of conservation and there is almost no information about the extinction risk of TSAR species on the IUCN Red List of threatened species. Nevertheless, we take this with strong caution for both practical and technical reasons. The TSAR group is home to ecologically important species such as *Macrocystis pyrifera* (giant kelp), but also to *Plasmodium falciparum*, the malaria parasite, which is a target for extinction rather than conservation. On the technical side, we have very little data informing ED estimates in TSAR. There are just a few phylogenetic studies informing the relationships of these taxa, and only 69 out of 48963 nodes have date estimates. We also lack phylogenetic information for 88.3% of nodes in that clade meaning our results for those nodes would be informed by taxonomy only. The high uncertainty in dates and relationships for this clade could be elevating PD scores.

With the development of new sequencing techniques, including higher-throughput methods, new species are now being rapidly added to the tree of life^48^ and it seems likely that older and less diverse clades will continue to be updated in terms of both species and topology as new data comes in. If these clades are understudied they may be updated to a greater degree than other clades that are already rich in described species. Future work should model the effects of missing undescribed species on the ED distributions of such clades because they would be expected to strongly affect ED scores of known species and PD to species richness ratios in understudied parts of the tree where these values are more extreme. In contrast ED scores are likely quite fixed in areas of the tree of life where almost all species are already described and in the dataset (such as mammals and birds). The large quantitative mismatch between TSAR, mammals and birds, may also be partly a symptom of limitations in the concept of PD at the scale of the complete tree of life. The very long branch lengths involved may not be directly proportional to the number of new features acquired (as PD assumes) and furthermore long branches provide more opportunities for ancestral features to be lost, which is also not taken into account by PD. An alternative in the form of Evolutionary Heritage (EvoHeritage) would be a fruitful direction for future work to address these issues^49^. Consideration of TSAR in the context of our study mostly highlights interesting future work to overcome data limitations in TSAR and elsewhere, as well as an opportunity to more deeply explore the limitations of PD as a universal concept across the complete tree of life.

The main finding of our study is our estimate for the total phylogenetic diversity of all life and ED values for all described species. As phylogenies capturing relationships and date estimates continue to be added to the Open Tree database the calculations produced by our method will be incrementally improved. We hope that this work will inspire the generation of a more automated pipeline for large scale PD calculations that will be broadly applicable in the future studies of biodiversity and streamline conservation biology research and practice.

### Material and methods

The methods consist of 1) data synthesis 2) calculating ED scores 3) calculating total unique PD and threatened PD of clades 4) calculating EDGE scores for species with IUCN Red List assessments 5) exploration of the results.

#### Data synthesis

We made use of two main data sources in our method. The first, from OneZoom^28,29^, consists of a tree topology for the complete tree of life, already aligned to IUCN Red List data. Second, a JSON data file giving dates for a subset of ancestral nodes identifiable in the Open Tree of life.

For context, the data tables from the OneZoom tree of life explorer^28,29^ were generated (by previous work^28^) by synthesising a curated backbone and other phylogenetic sources. The data consists of two data tables one with entries corresponding to nodes of the tree and the other with entries corresponding to leaves. In the nodes table, each row has a node ID and associated information including a parent field to give the node ID of that node’s parent node. The leaf_left and leaf_right fields enable rapid access to the set of the descendants in the leaf table^28^. In the leaves table each leaf (species) has a field giving its ‘Open Tree Taxonomy’ (OTT) ID, which is a unique id assigned by OpenTree to taxonomically named nodes as a canonical taxon identifier^50^. Both the leaves table and nodes table have a real_parent field, which allows us to identify the parent node of this node or leaf in the original data source. These “real_parent” identifiers are node ids in the OneZoom Tree of Life, which are not stable, but are always mapped wherever possible to node IDs in the Open tree of life synthetic tree. In contrast the “parent” field may point to a node generated only through a process of random polytomy splitting done as part of the synthesis described in^28,29^.

The JSON formatted dates were originally generated (by McTavish et al.^30^) from phylogenetic estimates of dates for internal nodes in published studies. These dates are mapped from published phylogenies to internal nodes of the OpenTree Synthetic tree using the OpenTree phylogenetic conflict APIs^31^ (https://github.com/OpenTreeOfLife/chronosynth), an extension of the concepts published in DateLife^51^. In these data, some dates correspond to nodes with their own OTT taxonomic ID whilst others are mapped to unnamed nodes in the Open Tree synthetic tree. These unnamed nodes are given an identifier based on the identifiers of two leaves for which that node is the most recent common ancestor. The full versioned set of dates for this analysis, including the published source tree for each date estimate, is available in the data depository.

We linked the nodes table from OneZoom^28^ with the JSON data from Open Tree of Life^30^ into a single table with node dates wherever known. In doing this node dates are only given to real nodes (i.e. not to nodes that only exist because of random polytomy resolution) (see figure 5). Considering that the OneZoom dataset also contains some node date estimates, we keep these date estimates and add the estimates provided by the JSON data from. Among all the 48013 dated nodes we used in our estimation, 33687 were provided by the JSON data from Open Tree of Life and 14326 were provided by OneZoom dataset. This process can assign multiple node date estimates to one node ID. Our strategy is to keep all of these date estimates and randomly choose one of them in each repeat for the following estimation of ED and PD scores. This jackknifing procedure gives us an estimate of uncertainty caused by selecting from inconsistent source data.

**Figure 5.**
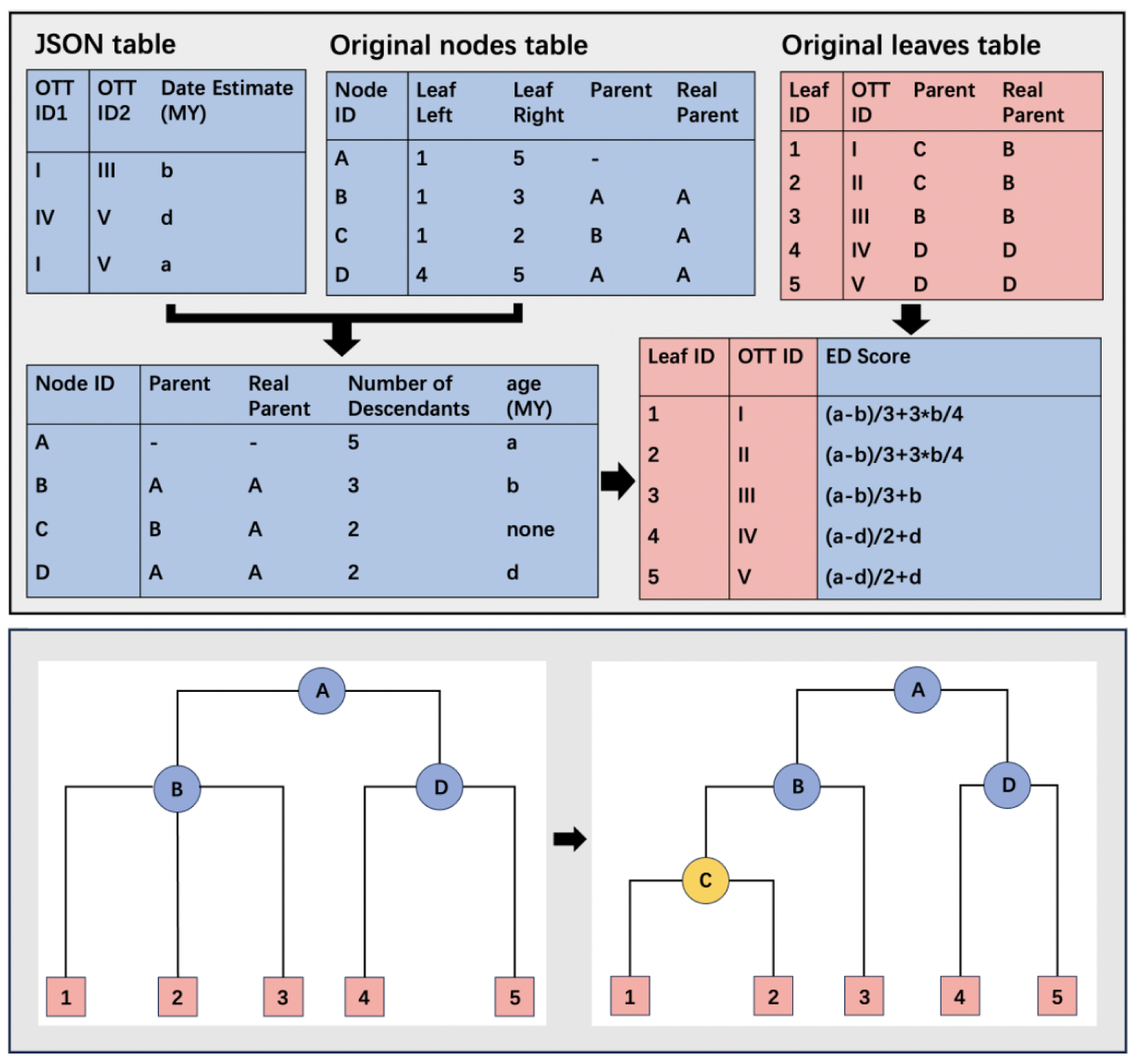
Overview schematic with an example tree showing the data we used and the process of calculating ED scores. Node IDs are the ones in the OneZoom database and are not stable but enable a neat structure for the tree being analyzed here. OTT taxon IDs are stable to updates in the tree data so are also included in our data tables (shown as roman numerals to differentiate them from the table IDs) and are used for most purposes. All leaves in the leaf table will be connected to a corresponding node in the node table. By finding date estimates and number of descendants for each parent from the re-assembled nodes table, the ED score for each leaf can be calculated.

#### ED scores

To calculate ED, each branch length is weighted by the number of species which are descended from it. The ED score for a species is equal to the total of all the weighted lengths of the branches on the path from the root of the tree to this species^10^. The ED of species *i*, written *ED*_i_, can thus be calculated by the following formula:

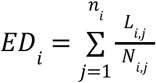

Where n_i_ gives the total number of ancestral branches to species *i*, *L*_*i*,1_ refers to the terminal branch length of species *i*, *L*_*i*,*j*_ for 2 ≤ *j* ≤ *n*_*i*_ gives the total length of interior branch *j* ancestral to species *i* and *N*_*i*,*j*_ gives the number of descendants for each of these same branches. The length of each branch can be calculated by the differences between two adjacent date estimates of nodes and the number of descendants for a node can be calculated by 1+ leaf_right - leaf_left (see ref^28^). In the calculation of ED scores for the entire tree of life, one major problem is that not all the nodes have a date estimate (For example, the node C in Figure 5). For the node without a date estimate, the branch length was estimated by calculating the total length between the closest ancestral node and the closest descendent node that have date estimates, and splitting that total length equally between all the sections created by undated nodes lying on the path between those two dated nodes. For instance, when calculating ED score for leaf 1 in Figure 5, the length of the branch connecting node B and node C is equal to the length of the branch that connects node C to leaf 1. This can be represented as b/2. Using this method the estimated ED score of leaf 1 (or leaf 2) can be calculated as:

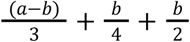

In order to capture a measure of uncertainty based on data deficiency, we conducted jackknife analysis by randomly removing 50% of node date estimates and using the rest of the data to perform all the analyses - repeating the process 1000 times. For the nodes that have multiple conflicting date estimates, we will randomly select one of the conflicting date estimates during the bootstrap analysis and discard the other(s). All the code used for the calculation of ED are available on GitHub (https://github.com/AlexGuojl/Phylogenetic-diversity-across-the-complete-tree-of-life).

##### Unique PD and threatened PD

Total unique PD of a given clade can be represented as the sum of the branches within this clade. With all the ED scores calculated, the total unique PD of a clade in the tree of life can be calculated by the sum of all ED scores minus the ED scores contributed by the branches between the root of the clade and the origin of life. The small deduction is necessary because the ED scores were all calculated in the context of the whole tree of life and so will consider every branch back to the origin even if contributions are negligible. For example, the total unique PD of clade C in figure 7 can be represented as:

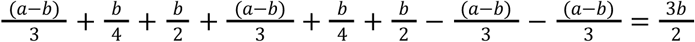

We use the ED scores generated in the 1000 jackknife replicates to calculate total unique PD for the monophyletic clades selected in this study. Each clade therefore has 1000 jackknife-based values of total unique PD. We produced box plots to see the distribution of total unique PD among the selected clades and all the PD scores were log transformed. By dropping all threatened species (VU, EN and CR species) from a given tree and subtracting the resulting PD from the original full PD, we estimated the threatened PD of monophyetic clades. We also incorporated a “worst case” situation in which the Data Deficient (DD) species and those species whose conservation status has not been evaluated (NE) by IUCN are regarded as threatened and therefore are dropped from the tree as well.

#### EDGE species

As a critical indicator of the health of the world’s biodiversity, the IUCN Red List of threatened species gives a categorical risk of extinction for successfully evaluated species^32,52^. By combining the calculated ED scores with the red list, we calculated EDGE scores using following formula set out in previous work^10^:

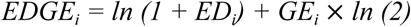

Where GE*_i_* is the extinction risk of species *i*, based on the IUCN red list (Least Concern=0, Near Threatened and Conservation Dependent = 1, Vulnerable=2, Endangered=3, Critically Endangered=4). All species with a GE score were sorted by their EDGE scores in descending order, both the top 20 (main text) and top 100 (supplementary table S1) EDGE species were presented.

To quantify the contribution of each node to the ED of these top EDGE species, we further calculated a cumulative ED value for each interior node, which is equal to the total weighted lengths of the branches on the path from the root to this node. The branch lengths are weighted in the same way as when calculating ED scores for a certain species. For example, in figure 5, the cumulative ED value of node B can be represented as:

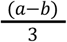

#### Exploration of the results

The ED score reported for each species is the median ED score among the 1000 ED scores generated in the jackknife analysis. The ED scores were log transformed to the base 10. To explore the distribution of ED scores across all described life, we divided all monophyletic clades into 6 groups based on their log transformed species richness (group 1: 1-2, group 2: 2-3, group 3: 3-4, group 4: 4-5, group 5: 5-6, group 6: above 6). Then we randomly sampled 4 clades within each group and made density plots of ED scores. We also arranged these clades into a phylogenetic tree based on their relationships in the OneZoom tree of life (see Figure S9).

For the purpose of further descriptive discussions of ED distributions within monophyletic clades we selected the following monophyletic groups: Telonemia, Stramenopile, Alveolates and Rhizaria (TSAR), Diaphoretickes, Spermatophyta, Chloroplastida, Holomycota, Metazoa, Mollusca, Chelicerata, Hymenoptera, Coleoptera, Diptera, Lepidoptera and Vertebrata. We further separated the Vertebrata clade into 9 clades: Cyclostomata, Chondrichthyes, Actinopterygii, Amphibia, Crocodylia, Testudines, Squamata, Mammalia and Aves. For each clade, we also compared the ED score of each species within this clade to the 0.99 quantile, 0.95 quantile and median value of the ED scores across the complete tree of life. We produced a stacked bar plot to see the proportion of high ED species across these clades (Figure 2). We devised a randomisation test to assess whether a clade disproportionately represents species that would be considered high ED (99 percentile rank) in the context of the complete tree of life. The number of species in the group (n) and the number of species in the group with ED ≥ Q(0.99) in the context of the complete tree (k) were recorded. We then performed 1000 iterations of randomly sampling n species from the entire "Biota" pool and counted, for each sample, how many of the species were in the 99th percentile of ED scores among all species in the complete tree of life. We then calculated the proportion of random samples where the number of top-ED species was ≥ k and ii) “proportion high”: the proportion of random samples where the number of top-ED species was < k. These two values provide a two-tailed interpretation: If this proportion was < 0.05 the group has significantly more high-ED species than expected (marked as "+"* in the plots). Conversely if this proportion was > 0.95, the group has significantly fewer high-ED species than expected (marked as "−"*). We visualized ED distributions across clades using violin plots combined with jittered points colored by quantile categories. Asterisks representing significance (*+, *-) were placed above each clade based on the comparison to the null distribution. This part of the analysis was conducted through R-4.4.2. To test the consistency of the ED scores estimated in this study with those of previous work, we compared our results against published data of ED and EDGE scores. These were 2020 values calculated via the original EDGE method for comparison with our results (Rikki Gumbs personal communication), not those using EDGE2 which would not be comparable to our values. We produced scatter plots to characterize the deviation between our values and the previously calculated values for all those species provided and calculated the mean percentage error on those scatter plots.

## Acknowledgements

The authors thank Rikki Gumbs for providing his estimation of ED and EDGE scores of selected monophyletic clades and his useful feedback on the manuscript. JR thanks the Leverhulme Trust for supporting his contribution to this work through a Research Fellowship (RF-2022-497).

The silhouette images of organisms used in Figure 1 and Figure S1 were collected from Phylopic (http://phylopic.org/). *Yochelcionella Cyrano*: Adapted from an illustration of Yochelcionella cyrano by Stanton F. Fink (vectorized by T. Michael Keesey). Licensed under CC BY-SA 3.0., *Helix aspersa*: Adapted from an illustration of Helix aspersa by Gareth Monger. Licensed under CC BY 3.0.. *Andrena barbara*: Photo by Sam Droege, vectorized by T. Michael Keesey, licensed under Creative Commons Attribution 3.0 Unported (CC BY 3.0). https://creativecommons.org/licenses/by/3.0/. *Gyrinus convexiusculus*: Adapted from an illustration by Desmond W. Helmore, uploaded by T. Michael Keesey. Licensed under CC BY 4.0., Micropezidae: Image by T. Michael Keesey, licensed under Creative Commons Attribution 3.0 Unported (CC BY 3.0). https://creativecommons.org/licenses/by/3.0/, *Micropezidae*: Adapted from a vector illustration by T. Michael Keesey, based on original work by Stanton F. Fink. Licensed under CC BY 3.0., *Ectoedemia coscoja*: Adapted from original work by E. J. Van Nieukerken, A. Laštůvka, and Z. Laštůvka (vectorized by T. Michael Keesey). Licensed under CC BY 3.0.,Macropus: Adapted image: Based on a photograph by Mathew Callaghan. Licensed under CC0 1.0 Universal Public Domain Dedication, *Balanerpeton woodi*: Adapted from an illustration by Scott Hartman. Licensed under CC BY 3.0. *Loligo vulgaris* by Cagri Cevrim, *Mesolimulus walchi* by Dean Schnabel, *Atrax robustus* by Margot Michaud, *Heterometrus laoticus* by Margot Michaud, *Martialis heureka* by Martialis heureka, *Lucanus* by Margot Michaud, *Carabidae* by Michael Day, *Toxomerus politus* by Andy Wilson, *Biston betularia* by Jamie Whitehouse, *Papilio polyxenes* by Andy Wilson, *Apteryx australis* by Steven Traver, *Petromyzon marinus* by Christoph Schomburg, *Oncorhynchus nerka* by xgirouxb, *Agalychnis lemur* by Margot Michaud, *Crocodylus acutus* by Beth Reinke, *Uroplatus phantasticus* by Steven Traver, *Tachyglossus aculeatus* by Becky Barnes, *Menura* by T. Michael Keesey are in the public domain (CC0 1.0). No attribution required.

The images used in Figure 6 are credited as follows: were collected from different resource: L. chalumnae: Image by Bruce A.S. Henderson, CC BY 4.0, via Wikimedia Commons. N. fosteri: Image by Kenneth Lu, CC BY 4.0., via Inaturalist, M. halope: Peter Parks, public domain, via Wikimedia Commons., E.madagascariensis (Madagascar Bighead Turtle). Photo by Tony King, CC BY 4.0., via Inaturalist. D.mawii: Photo by Giovana A. Valencia, licensed under CC BY 4.0. via Inaturalist. L. archeyi: photographed by Carey_Knox_Southern_Scales. L.nigriventer: Modified from original. Photo by Uzi Paz, via Pikiwiki Israel. Licensed under CC BY 2.5., P.megacephalum: Lawrence Hylton, licensed under CC BY 4.0, via Inaturalist.; M.chejuensis: Photograph by Kim, Hyun-tae, sourced from iNaturalist, licensed under Creative Commons Attribution 4.0 International (CC BY 4.0).; A.sturio: Photograph by Aah-Yeah, sourced from Flickr, licensed under Creative Commons Attribution 2.0 Generic (CC BY 2.0); M.calocoma: Photo by Eric Knight, licensed under CC BY. C.lamottei: Endemic amphibians of Mt Oku. Image from Doherty-Bone TM, Gvoždík V (2017), The Amphibians of Mount Oku, Cameroon: an updated species inventory and conservation review, ZooKeys 643: 19–139. https://doi.org/10.3897/zookeys.643.9422. Licensed under CC BY 4.0.

## Supplementary materials

**Figure S1.**
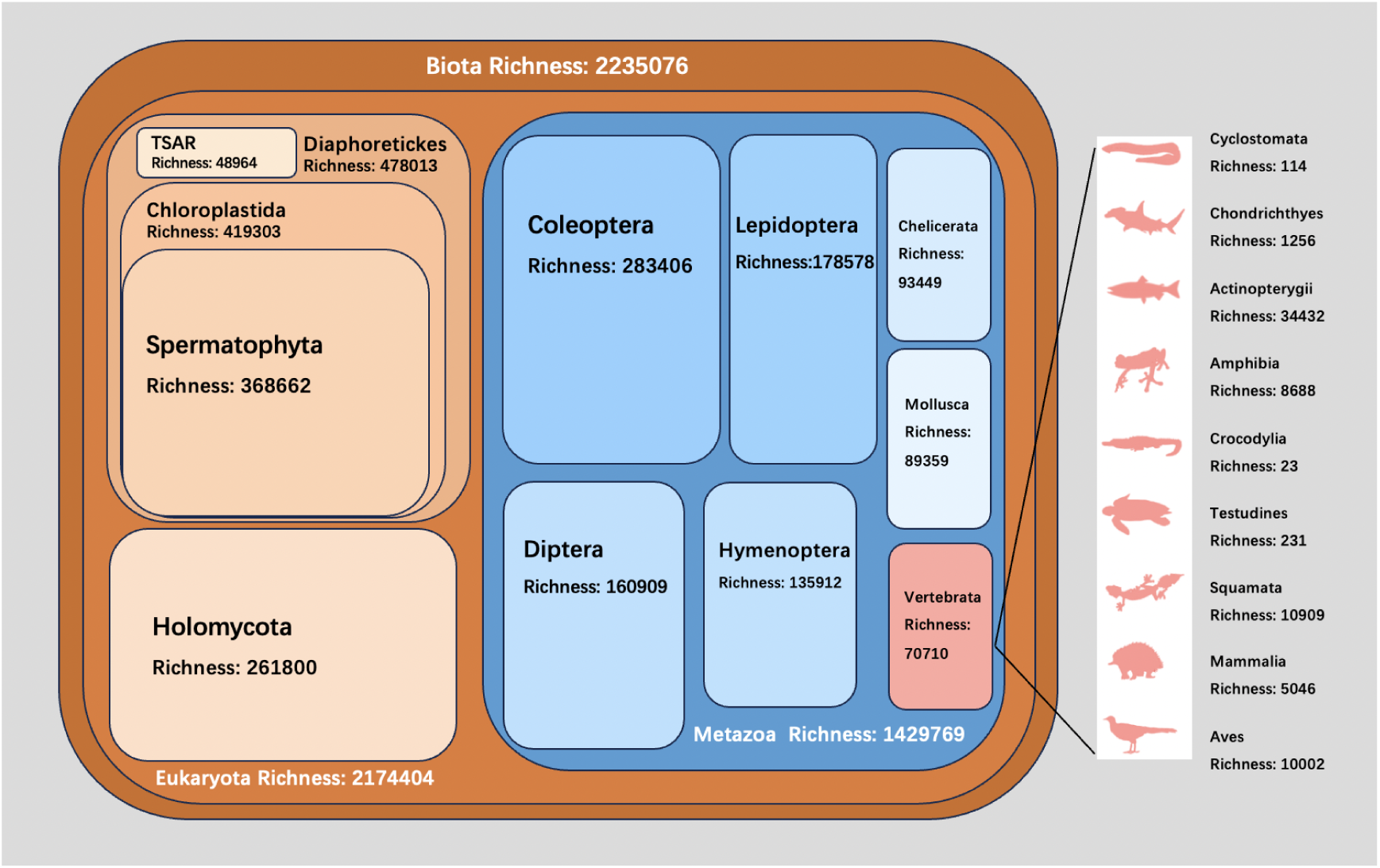
The relationship between the selected monophyletic clades for this study. In this figure we illustrate all our choices of taxa and the hierarchical relationships between them. The size of each rectangle and the depth of its color qualitatively reflect the species richness of each clade to help with readability.

**Figure S2.**
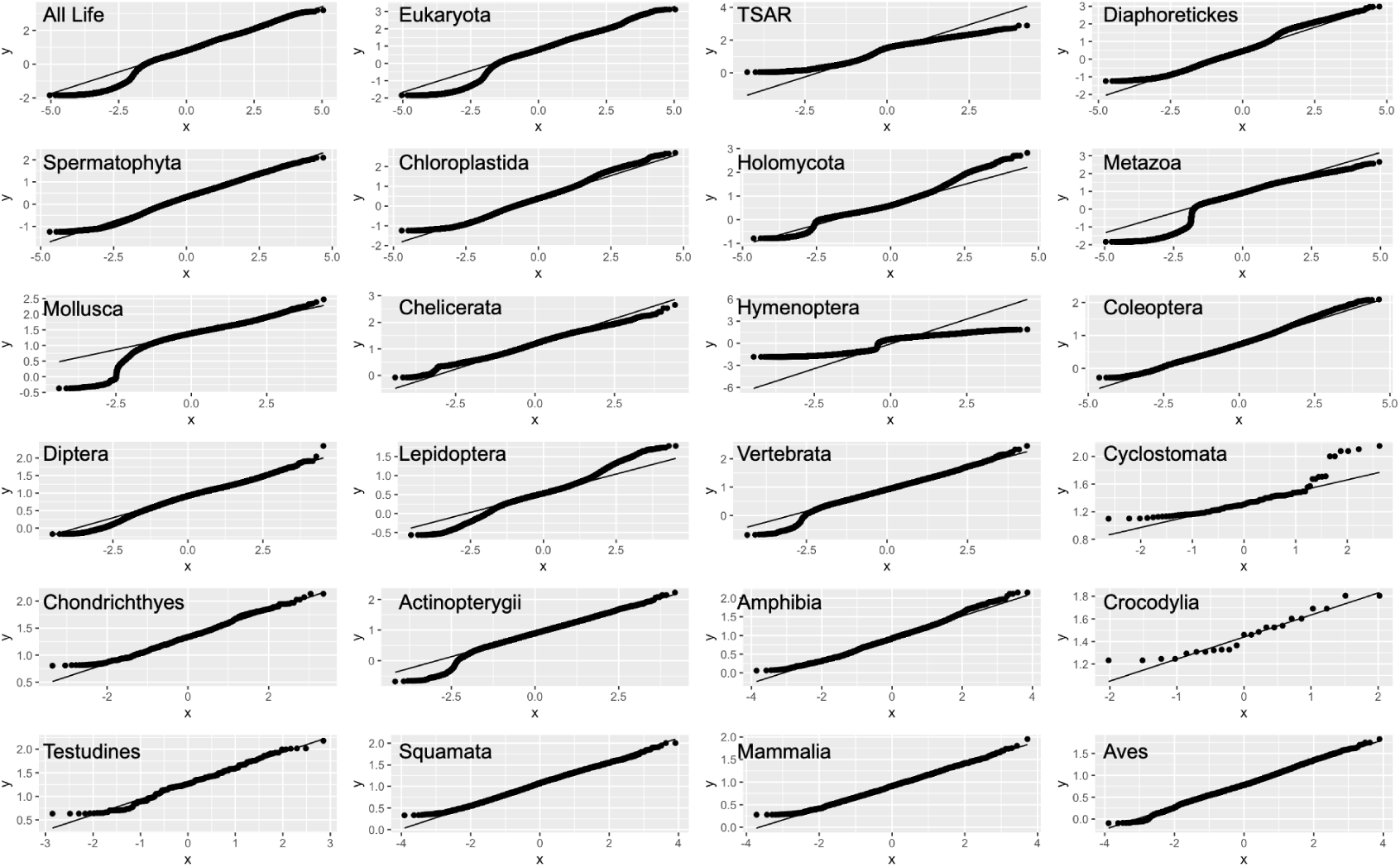
Q-Q plot of ED scores for our 24 selected groups. All of the selected groups have points that deviate from the straight diagonal line. This indicates that none of those groups has log-normally distributed ED scores.

**Figure S3.**
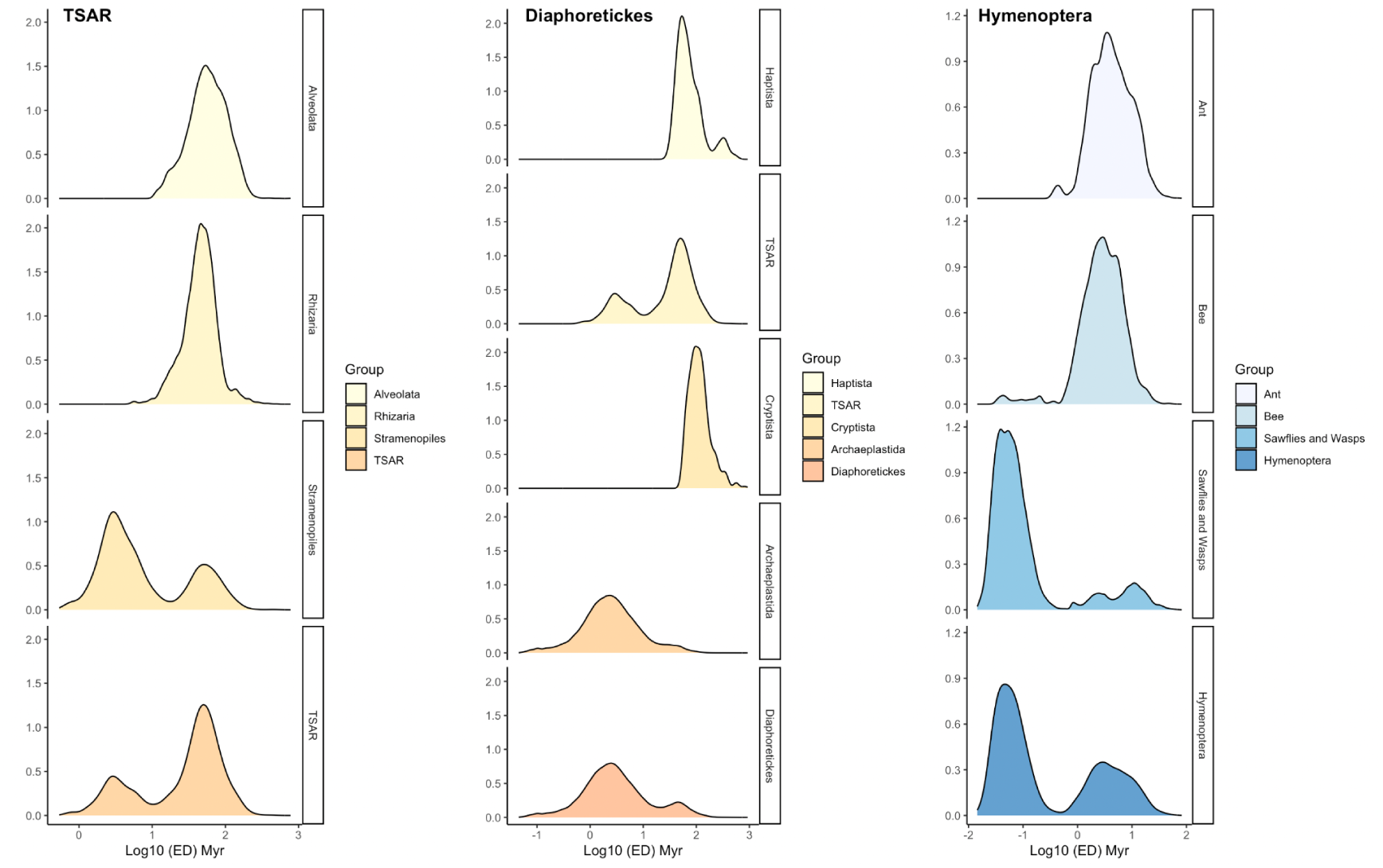
the distribution of ED scores in Eukaryota subgroups. The selected Eukaryota groups which show bimodal distribution of ED scores were further divided into subgroups. The depth of the color refers to the species richness of each subgroup.

**Figure S4.**
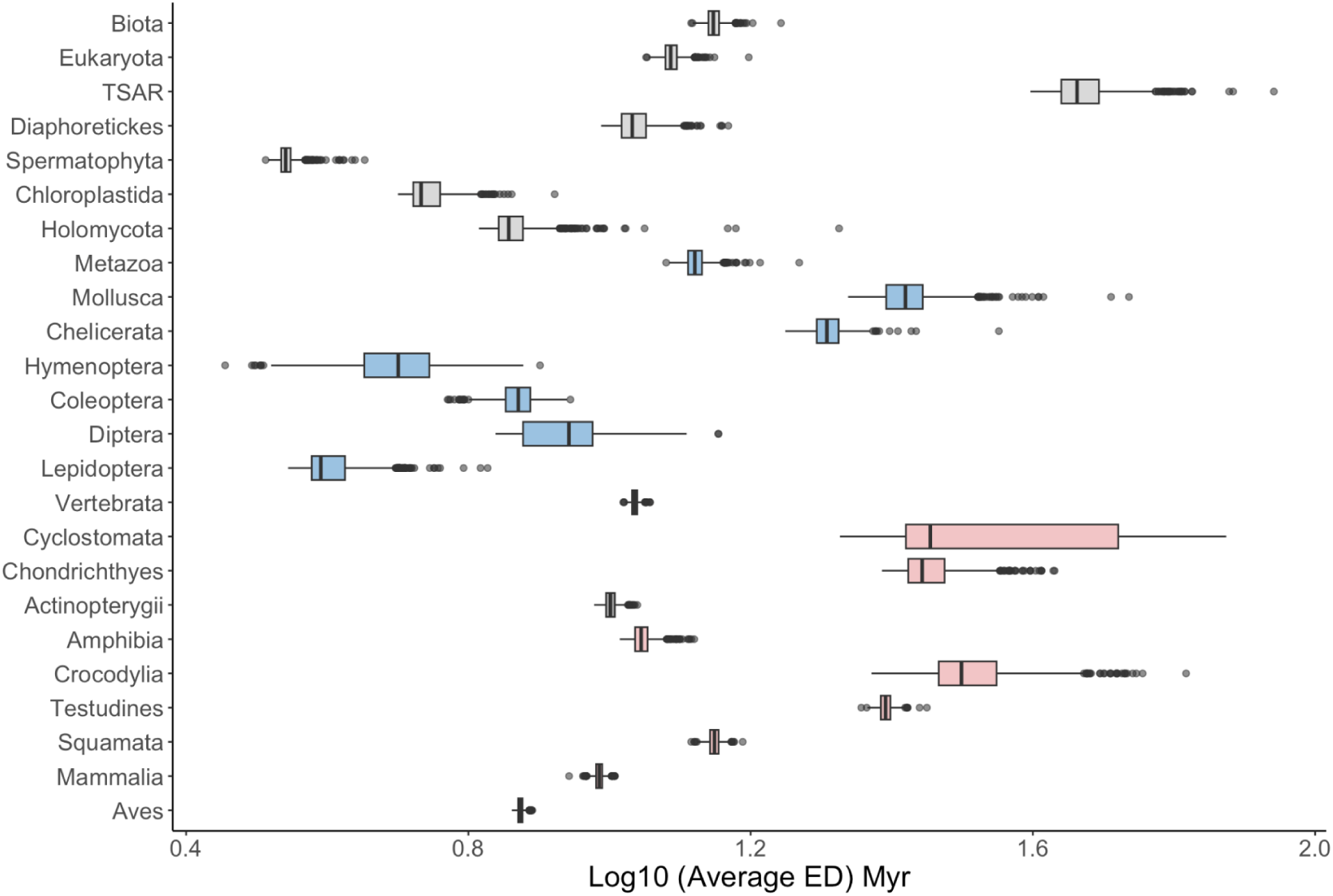
The average ED scores for the selected clades. The average values are calculated by summing the ED scores of its corresponding group and divided by the species richness of that group. Each group has 1000 average ED scores aring from the jackknife analysis.

**Figure S5.**
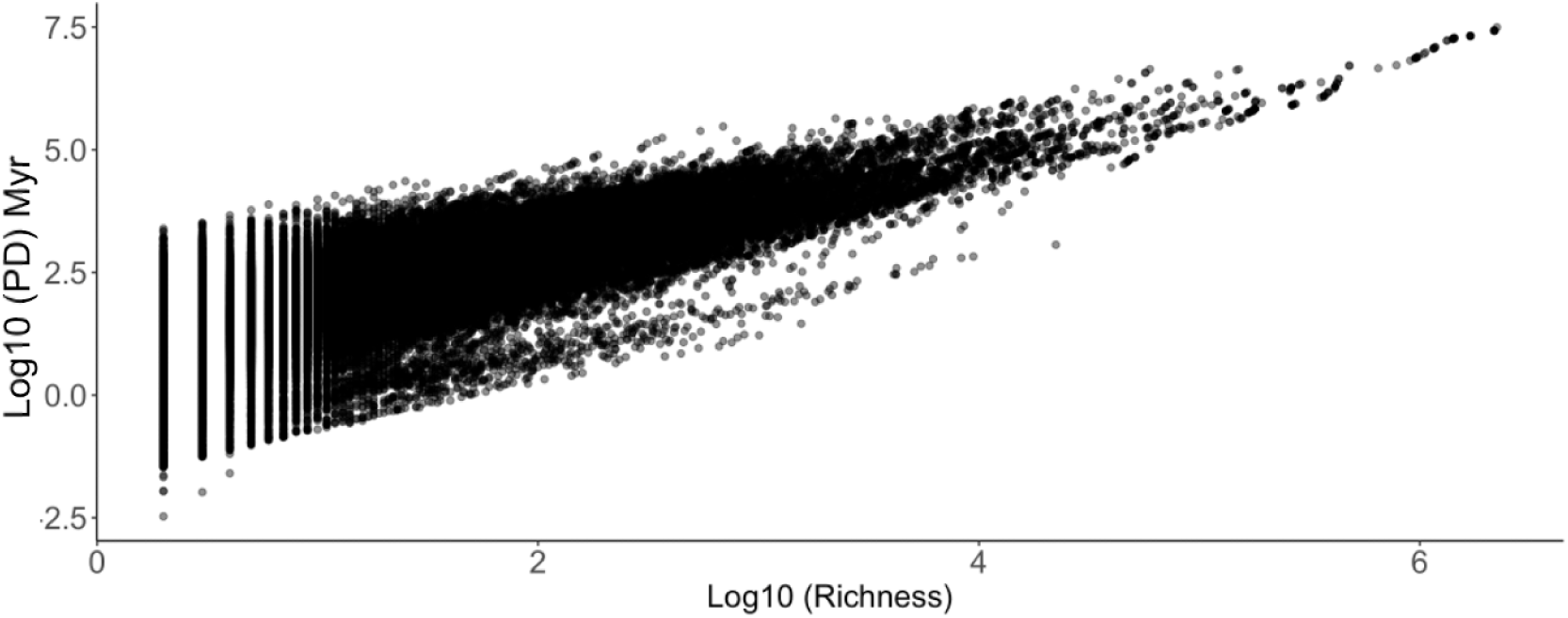
PD and species richness comparison. To test the relationship between total unique PD and species richness for all clades, we made a scatter plot to show the relationship between the log median PD among the 1000 repeats and log species richness. Our results for all clades showed that the total unique PD increases approximately according to a power law with the species richness. (figure 34. Rs = 0.729, p ≪ 0.0001;) although there is substantive variability especially for smaller species richness values.

**Figure S6.**
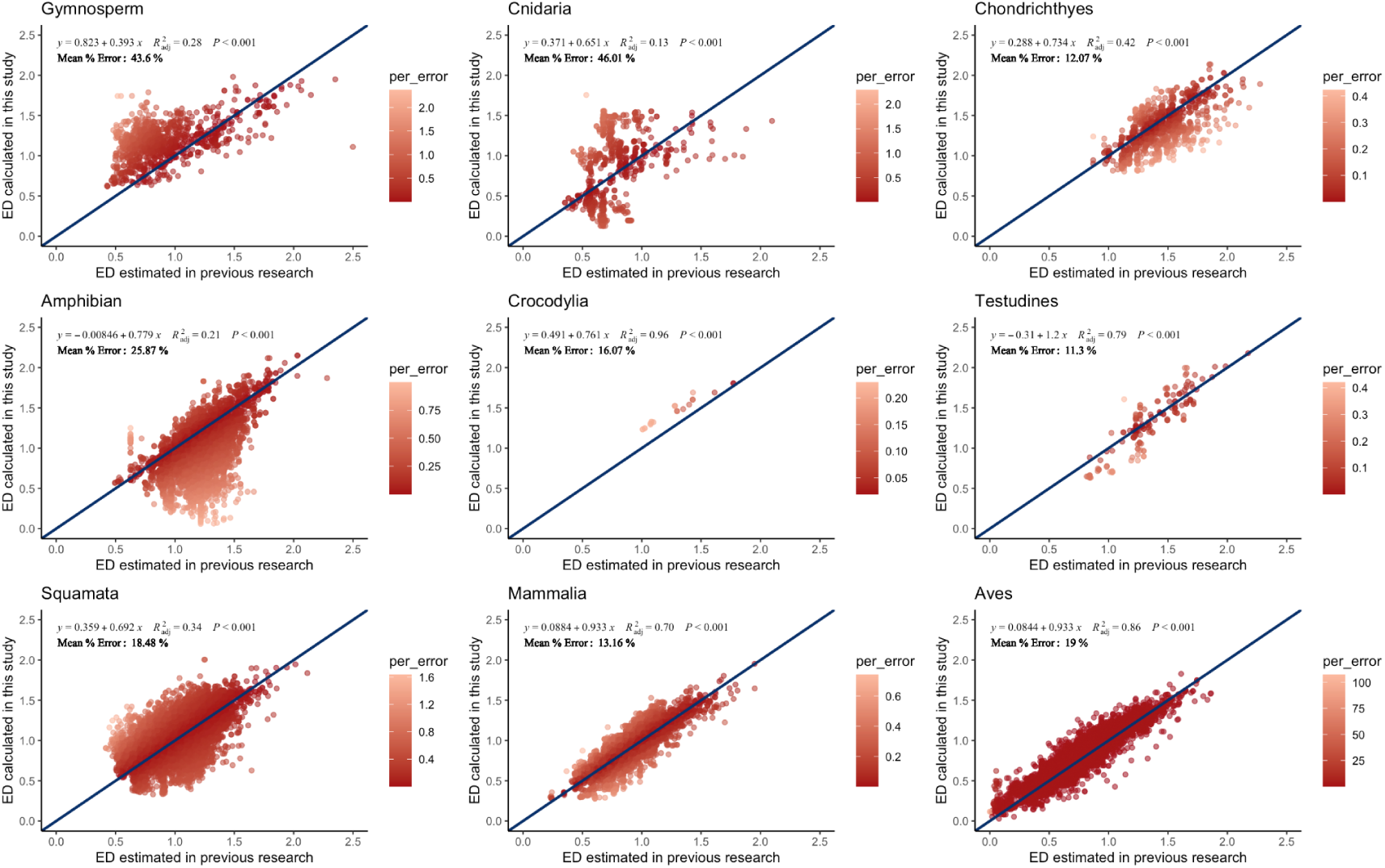
Comparison of ED scores. The x-axis shows the existing published values (Gumbs et al. 2018), and the y-axis is the values calculated in our study. The diagonal line is shown for illustrative purposes with an intercept of 0 and a slope of 1.

**Figure S7.**
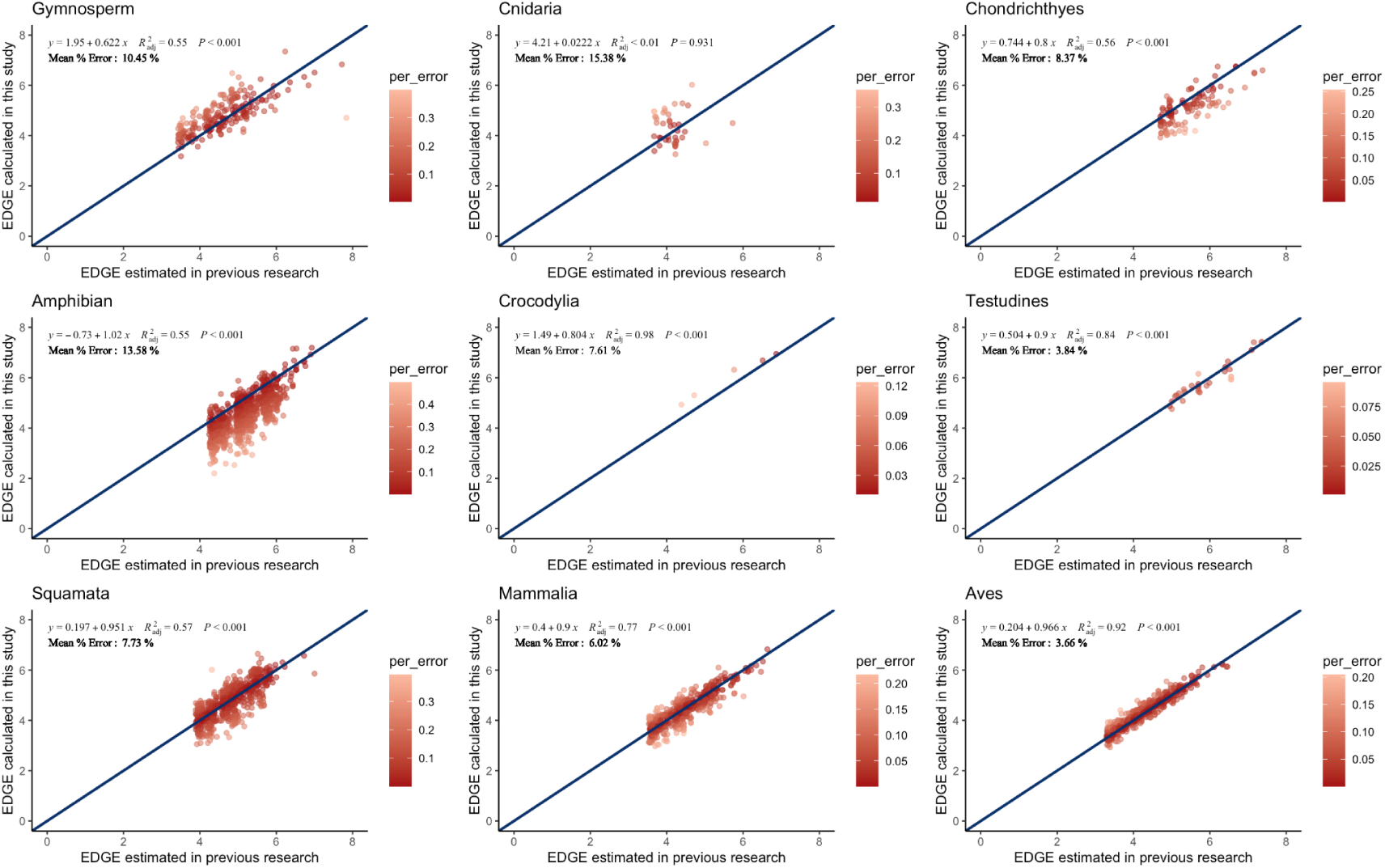
Comparison of EDGE scores. The x-axis shows the existing published values (Gumbs et al. 2018), and the y-axis is the values calculated in our study. The diagonal line is shown for illustrative purposes with an intercept of 0 and a slope of 1.

**Figure S8.**
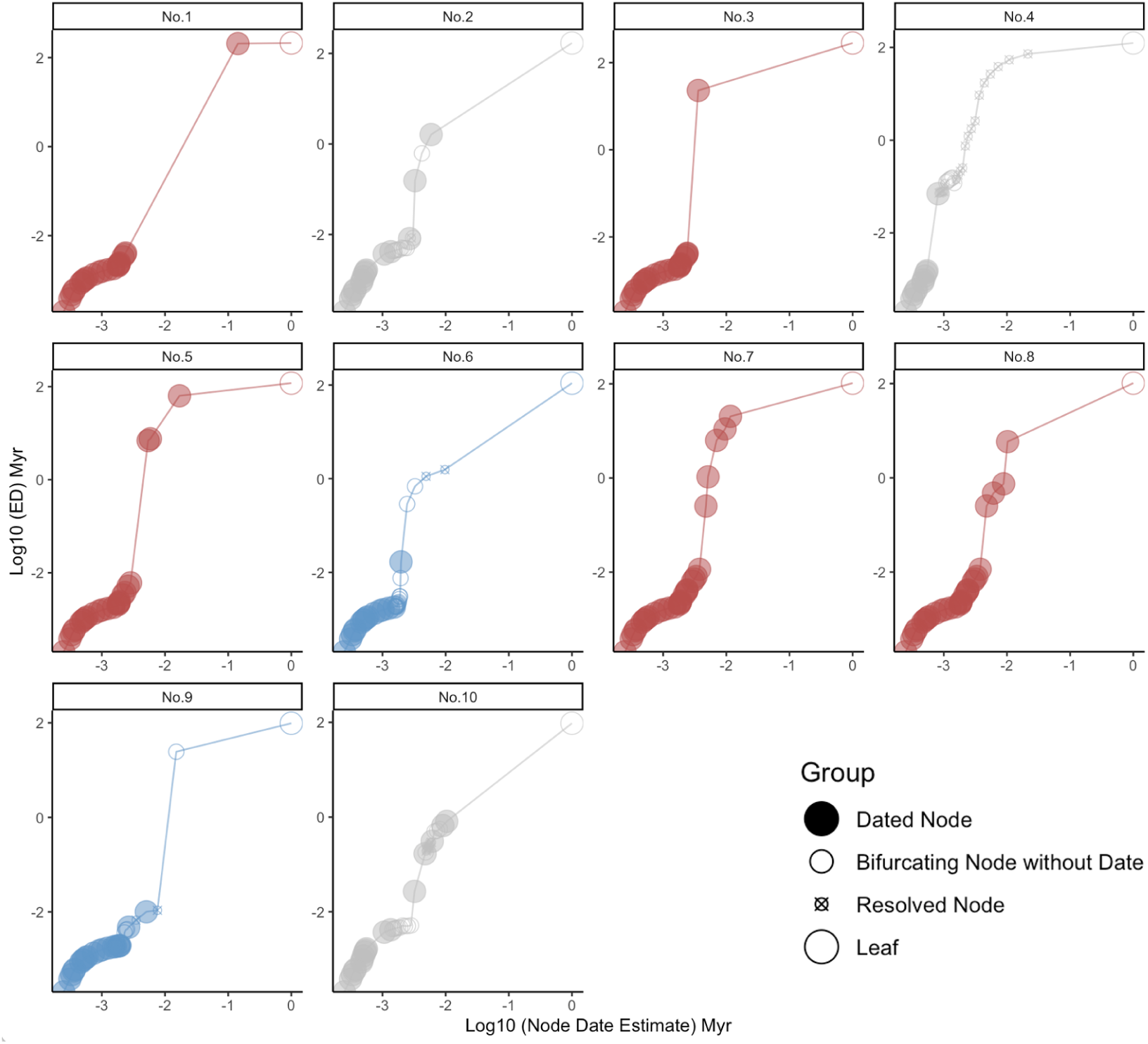
Contribution of different nodes to the ED score of top 10 EDGE species. The different colors of points correspond to different species, the shape and size of different points indicates the type of this node. “Dated nodes” have the strongest quantitative backing with both a date and a topology known in the source data. “Bifurcating nodes” have a known topology without polytomy, the date was only interpolated based on other dated nodes. “Resolved nodes” appear in the data only through random polytomy resolution.

**Figure S9.**
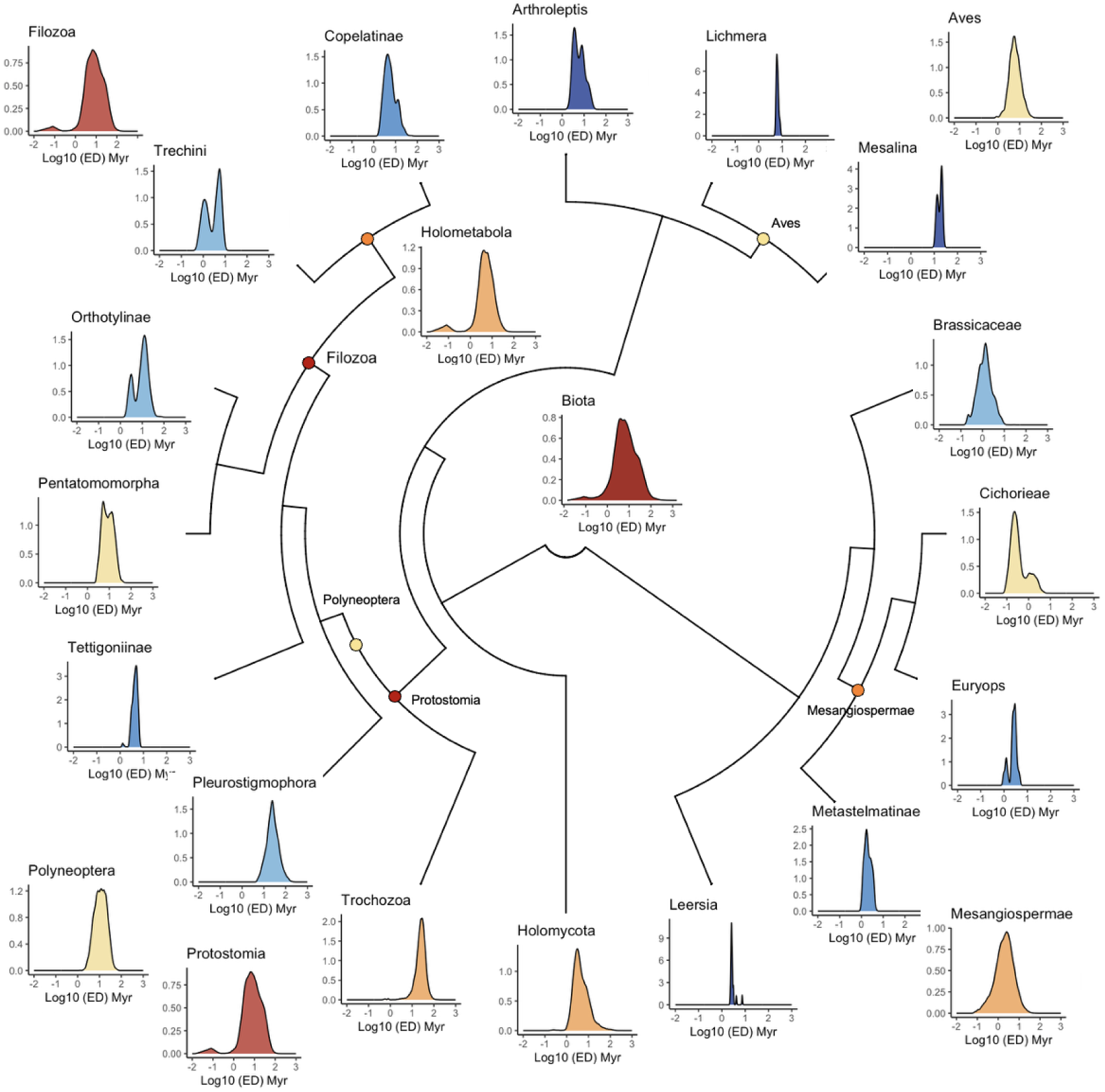
Distribution of ED scores for our selected clades showing their positions of the phylogeny.

**Supplementary data S1**

Median ED scores for all species in our dataset See https://zenodo.org/records/15688295

**Supplementary data S2**

Unique PD across the tree of life

See https://zenodo.org/records/15688492

**Supplementary data S3**

Database dumps provided by OneZoom with leaves and nodes information See https://zenodo.org/records/16810118

**Supplementary data S4**

Citations and quantitative comparisons between our work and previous work. Covers PD and mean ED results for selected clades where data for comparison is available.

See the supplementary information file attached.

**Supplementary table S1.** The top 100 EDGE species.

| Rank | Name | ED (Myr) | EDGE | Conservation Status | 95% Confidence Interval of ED (Myr) |
| --- | --- | --- | --- | --- | --- |
| 1 | *Latimeria chalumnae* | 213.02 | 8.14 | CR | 199.90-210.32 |
| 2 | *Mankya chejuensis* | 170.09 | 7.91 | CR | 132.53-137.27 |
| 3 | *Neoceratodus forsteri* | 282.61 | 7.73 | EN | 236.47-244.69 |
| 4 | *Galaxaura barbata* | 124.51 | 7.6 | CR | 140.64-144.07 |
| 5 | *Acipenser sturio* | 118.78 | 7.56 | CR | 115.49-117.18 |
| 6 | *Acharax alinae* | 109.66 | 7.48 | CR | 74.11-78.41 |
| 7 | *Erymnochelys madagascariensis* | 103.73 | 7.42 | CR | 95.68-97.10 |
| 8 | *Dermatemys mawii* | 102.49 | 7.41 | CR | 88.61-90.63 |
| 9 | *Mictocaris halope* | 98.01 | 7.37 | CR | 94.14-95.96 |
| 10 | *Microcycas calocoma* | 95.74 | 7.34 | CR | 67.66-71.57 |
| 11 | *Euastacus girurmulayn* | 93.32 | 7.32 | CR | 82.16-88.95 |
| 12 | *Eostemmiulus caecus* | 85.53 | 7.23 | CR | 85.09-85.37 |
| 13 | *Geothallus tuberosus* | 85.39 | 7.23 | CR | 81.30-82.36 |
| 14 | *Antrisocopia prehensilis* | 85.35 | 7.23 | CR | 83.17-84.95 |
| 15 | *Nanocopia minuta* | 82.51 | 7.2 | CR | 80.33-81.75 |
| 16 | *Leiopelma archeyi* | 81.90 | 7.19 | CR | 81.85-83.08 |
| 17 | *Symmetromphalus hageni* | 80.06 | 7.17 | CR | 79.55-80.53 |
| 18 | *Crotaphatrema lamottei* | 79.35 | 7.16 | CR | 77.30-78.57 |
| 19 | *Latonia nigriventer* | 78.63 | 7.15 | CR | 70.65-72.16 |
| 20 | *Platysternon*<br>*megacephalum* | 76.59 | 7.12 | CR | 62.49-64.35 |
| 21 | *Speleoperipatus spelaeus* | 76.17 | 7.12 | CR | 75.11-76.51 |
| 22 | *Carettochelys insculpta* | 151.26 | 7.11 | EN | 111.67-116.45 |
| 23 | *Planorbidella depressa* | 73.00 | 7.08 | CR | 72.05-72.98 |
| 24 | *Spelaeoecia bermudensis* | 72.36 | 7.07 | CR | 76.67-77.76 |
| 25 | *Huso huso* | 72.01 | 7.06 | CR | 73.32-75.90 |
| 26 | *Opisthopatus roseus* | 71.11 | 7.05 | CR | 69.25-70.04 |
| 27 | *Scaphirhynchus suttkusi* | 68.01 | 7.01 | CR | 67.70-70.43 |
| 28 | *Sphaeromimus lavasoa* | 66.68 | 6.99 | CR | 64.48-65.87 |
| 29 | *Hymenophyllum*<br>*helicoideum* | 66.58 | 6.99 | CR | 66.44-67.08 |
| 30 | *Acipenser stellatus* | 66.31 | 6.98 | CR | 63.12-64.61 |
| 31 | *Pseudoscaphirhynchus*<br>*kaufmanni* | 66.31 | 6.98 | CR | 63.15-64.62 |
| 32 | *Margaritifera marocana* | 65.83 | 6.97 | CR | 64.09-64.59 |
| 33 | *Phycolepidozia exigua* | 65.53 | 6.97 | CR | 63.95-64.78 |
| 34 | *Margaritifera hembeli* | 65.25 | 6.97 | CR | 64.97-65.84 |
| 35 | *Microsphaerotherium*<br>*anjozorobe* | 65.24 | 6.97 | CR | 70.06-71.23 |
| 36 | *Seychellia lodoiceae* | 65.12 | 6.96 | CR | 39.31-42.39 |
| 37 | *Pentaplebia gamblesi* | 64.31 | 6.95 | CR | 63.40-64.20 |
| 38 | *Gigantopelta aegis* | 64.24 | 6.95 | CR | 66.13-67.08 |
| 39 | *Acipenser schrenckii* | 63.85 | 6.94 | CR | 67.48-69.03 |
| 40 | *Alligator sinensis* | 63.83 | 6.94 | CR | 57.57-60.18 |
| 41 | *Isoetes hewitsonii* | 63.75 | 6.94 | CR | 64.44-66.30 |
| 42 | *Acipenser nudiventris* | 63.74 | 6.94 | CR | 66.59-68.24 |
| 43 | *Riotintobolus minutus* | 63.63 | 6.94 | CR | 63.10-63.93 |
| 44 | *Pleiodon ovatus* | 63.59 | 6.94 | CR | 63.40-64.01 |
| 45 | *Colossobolus pseudoaculeatus* | 63.21 | 6.93 | CR | 69.96-72.26 |
| 46 | *Isoetes sinensis* | 62.92 | 6.93 | CR | 67.41-69.45 |
| 47 | *Andrias davidianus* | 62.61 | 6.93 | CR | 64.52-65.29 |
| 48 | *Mahezomus apicoporus* | 62.25 | 6.92 | CR | 61.29-62.17 |
| 49 | *Procaris chacei* | 60.99 | 6.9 | CR | 55.62-58.06 |
| 50 | *Huso dauricus* | 60.77 | 6.9 | CR | 82.39-86.34 |
| 51 | *Aneuretus simoni* | 60.65 | 6.89 | CR | 59.12-59.65 |
| 52 | *Rhynchobatus luebberti* | 60.59 | 6.89 | CR | 62.71-64.08 |
| 53 | *Rhynchobatus immaculatus* | 60.59 | 6.89 | CR | 62.68-64.06 |
| 54 | *Euastacus yigara* | 60.56 | 6.89 | CR | 59.16-59.98 |
| 55 | *Vandiemena ratkowskiana* | 60.52 | 6.89 | CR | 51.80-54.18 |
| 56 | *Leptogyra inflata* | 60.40 | 6.89 | CR | 63.34-64.61 |
| 57 | *Perissolestes remus* | 60.32 | 6.89 | CR | 61.92-65.98 |
| 58 | *Acipenser dabryanus* | 60.05 | 6.88 | CR | 53.86-56.16 |
| 59 | *Acipenser sinensis* | 60.05 | 6.88 | CR | 53.84-56.20 |
| 60 | *Sphaeromimus saintelucei* | 59.68 | 6.88 | CR | 61.54-62.31 |
| 61 | *Spiromimus albipes* | 59.20 | 6.87 | CR | 58.48-58.98 |
| 62 | *Engaewa pseudoreducta* | 58.79 | 6.86 | CR | 59.73-63.27 |
| 63 | *Branchinecta belki* | 58.45 | 6.86 | CR | 58.57-59.26 |
| 64 | *Sternothyris braueri* | 116.47 | 6.85 | EN | 97.48-99.85 |
| 65 | *Garypus titanius* | 57.72 | 6.85 | CR | 56.67-57.42 |
| 66 | *Lirapex politus* | 57.69 | 6.84 | CR | 56.53-57.28 |
| 67 | *Sphaeromimus andrahomana* | 57.25 | 6.84 | CR | 57.44-58.03 |
| 68 | *Wollemia nobilis* | 56.91 | 6.83 | CR | 61.83-64.57 |
| 69 | *Zaglossus attenboroughi* | 56.79 | 6.83 | CR | 47.80-49.74 |
| 70 | *Galeorhinus galeus* | 56.77 | 6.83 | CR | 58.41-59.92 |
| 71 | *Lasmigona decorata* | 56.76 | 6.83 | CR | 59.99-62.53 |
| 72 | *Rhynchobatus australiae* | 56.56 | 6.83 | CR | 56.28-57.07 |
| 73 | *Margaritifera auricularia* | 56.50 | 6.82 | CR | 51.92-54.12 |
| 74 | *Acipenser mikadoi* | 56.48 | 6.82 | CR | 56.14-56.86 |
| 75 | *Alasmidonta raveneliana* | 56.30 | 6.82 | CR | 55.81-56.59 |
| 76 | *Solemya flava* | 55.32 | 6.8 | CR | 50.12-50.85 |
| 77 | *Rhombuniopsis tauriformis* | 55.25 | 6.8 | CR | 55.13-55.88 |
| 78 | *Atya brachyrhinus* | 55.11 | 6.8 | CR | 59.08-60.05 |
| 79 | *Granitobolus endemicus* | 54.97 | 6.8 | CR | 48.13-50.06 |
| 80 | *Zaglossus bruijini* | 54.47 | 6.79 | CR | 39.84-41.66 |
| 81 | *Milyeringa justitia* | 54.29 | 6.79 | CR | 58.24-60.66 |
| 82 | *Iheyaspira lequios* | 54.18 | 6.78 | CR | 48.63-50.42 |
| 83 | *Riotintobolus mandenensis* | 54.05 | 6.78 | CR | 56.36-57.15 |
| 84 | *Colossobolus litoralis* | 54.04 | 6.78 | CR | 54.23-54.80 |
| 85 | *Pinna nobilis* | 53.69 | 6.77 | CR | 55.56-56.60 |
| 86 | *Sphaeromimus splendidus* | 53.55 | 6.77 | CR | 55.61-56.60 |
| 87 | *Lamellomphalus manusensis* | 53.33 | 6.77 | CR | 45.79-49.61 |
| 88 | *Rhynchobatus laevis* | 53.21 | 6.77 | CR | 52.36-53.04 |
| 89 | *Rhynchobatus springeri* | 53.21 | 6.77 | CR | 55.20-55.91 |
| 90 | *Afrogarypus seychellesensis* | 53.16 | 6.76 | CR | 52.54-53.35 |
| 91 | *Branchinecta mexicana* | 53.10 | 6.76 | CR | 53.30-53.77 |
| 92 | *Latimeria menadoensis* | 213.02 | 6.75 | VU | 49.49-50.91 |
| 93 | *Squatina oculata* | 52.28 | 6.75 | CR | 49.96-51.58 |
| 94 | *Squatina squatina* | 52.28 | 6.75 | CR | 49.99-51.56 |
| 95 | *Squatina aculeata* | 52.28 | 6.75 | CR | 200.20-210.62 |
| 96 | *Colossobolus minor* | 52.02 | 6.74 | CR | 53.61-54.33 |
| 97 | *Hylekobolus andasibensis* | 52.01 | 6.74 | CR | 50.53-50.92 |
| 98 | *Rhynchobatus cooki* | 51.48 | 6.73 | CR | 53.13-54.58 |
| 99 | *Spiromimus litoralis* | 50.56 | 6.72 | CR | 51.59-52.20 |
| 100 | *Atractosteus tristoechus* | 50.52 | 6.71 | CR | 61.39-64.65 |

